# Impact of Coding Region Accessibility and Uridine Chemistry on Translation in mRNA–DNA Hybrid Origami

**DOI:** 10.64898/2026.06.25.734245

**Authors:** Carmine D’Amico, Miska Mykkänen, Sharon Saarinen, Ville Säkkinen, Mauri A. Kostiainen

## Abstract

Messenger RNA (mRNA) is a prerequisite for programmable protein expression, but its therapeutic and synthetic-biology applications are limited by instability and susceptibility to degradation. Hybridizing a mRNA with short DNA staples can fold it into a defined nanostructure and shield it from nuclease degradation, but the same base pairing hides the transcript from the ribosome. Which regions must remain accessible, and how much coding sequence can be given over to structure, is not known. We folded an EGFP-encoding mRNA into a six-helix bundle and varied which parts of the open reading frame were left unpaired, measuring output both within the folded series and against the unhybridized transcript. The requirement is positional and asymmetric: exposing the first 35 coding nucleotides raises translation, exposing the last 35 does not. More exposure is not better: across overhangs of 35 to 100 nucleotides output falls in two discrete steps, with no difference between 50, 60, 70 and 80 nucleotides, indicating a threshold set by the ribosomal initiation footprint rather than a graded dependence on length. A construct with the entire open reading frame paired still translates at two-thirds of the reference, implying that elongating ribosomes displace staples as they read and that the folded state is transient during translation. Importantly, one staple set folds scaffolds built from uridine, 5-methoxyuridine or N1-methylpseudouridine, and a 35-nucleotide overhang is optimal in all three. N1-methylpseudouridine gives the highest output but is threefold more sensitive to an over-long overhang, consistent with its stabilization of RNA secondary structure. Coding-overhang accessibility and uridine chemistry are therefore separable design parameters. Together these results begin to define how a folded mRNA can be made both stable and efficiently translated.

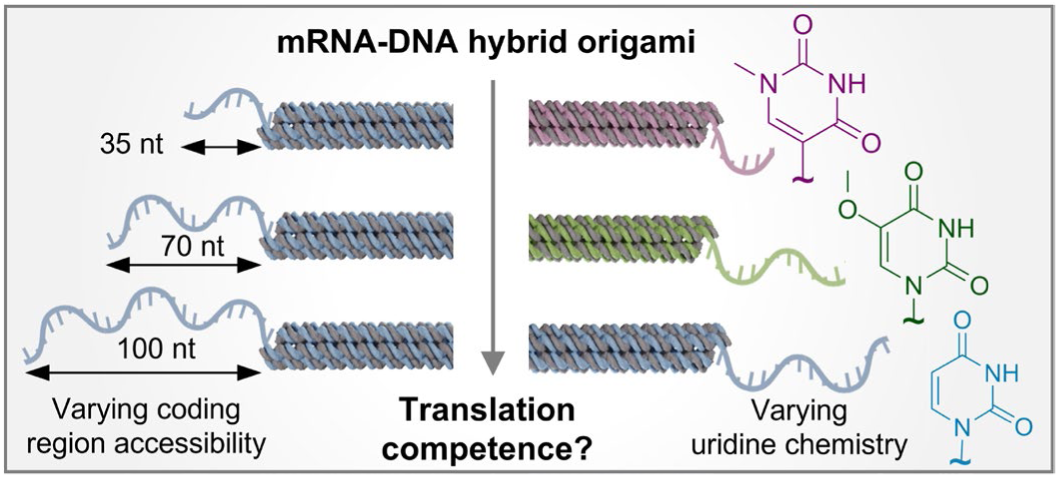

## Introduction

Synthetic messenger RNA (mRNA) has emerged as a versatile therapeutic platform, with applications spanning prophylactic vaccination, protein replacement, and gene editing support.^1^ The clinical validation of mRNA-based vaccines against SARS-CoV-2 demonstrated that in vitro-transcribed mRNA can direct protein synthesis at therapeutically relevant levels in humans, accelerating interest in extending the modality to more demanding indications.^2,3^ Nucleoside modifications have substantially improved immune evasion and translational output of synthetic mRNA,^4,5^ and systematic optimization of 5′ and 3′ untranslated regions has further increased expression levels and transcript stability.^6,7^ These advances operate at the level of sequence and chemistry, and they have been transformative. However, what they do not address is the three-dimensional structural state of the mRNA molecule itself.

Among the modifications available, the substitution of uridine has proven especially consequential. Pseudouridine, 5-methoxyuridine (5moU), and N1-methylpseudouridine (N1MeΨ) each reduce innate immune activation and alter translational output relative to unmodified uridine,^5^ and N1MeΨ in particular has become the clinical standard: both authorised SARS-CoV-2 mRNA vaccines replace uridine entirely with N1MeΨ, following the demonstration that it outperforms pseudouridine in protein expression while minimising immunogenicity.^8,9^ The benefit of these modifications is not solely immunological. In cell-free systems that lack innate immune sensing, N1-methylpseudouridine still enhances translation, increasing ribosome density and loading through effects on the ribosomal decoding centre that operate independently of immune evasion, while also reshaping the global secondary structure of the transcript.^10^ Different uridine analogues therefore differ both in how they fold and in how they are read by the ribosome. Yet how these modifications behave once the mRNA is constrained within a structured nanoparticle, where the chemistry of the nucleobase influences not only ribosomal decoding but also the duplex stability, has not been examined. Whether the translational hierarchy among uridine analogues established for linear mRNA is preserved, amplified, or overridden in highly structured systems is unknown, and is a question this work addresses directly.

Beyond sequence and chemistry, the structural state of the mRNA is itself a functional variable. Secondary structure is not a stochastic byproduct of sequence but a position-dependent feature whose deliberate design shapes protein output. For example, reduced structure in the 5′ leader and the first codons facilitates ribosomal scanning and initiation, while increased structure in the remainder of the coding sequence and the 3′ UTR protects the transcript and extends the period over which it is translated.^11^ Computational tools such as LinearDesign exploit this by jointly optimising secondary structure and codon usage to improve half-life and expression by orders of magnitude over codon optimisation alone.^12^ Yet these sequence-level design strategies leave the three-dimensional structural state of the folded mRNA molecule, and in particular its behaviour during and after encapsulation, largely unaddressed. Structure also governs the molecule’s physical behaviour during delivery: engineering rigid mRNA folding architectures amplifies protein expression several-fold in vivo relative to unstructured controls, through enhanced endosomal processing and prolonged intracellular retention.^13^ Also scattering studies of mRNA–lipid nanoparticle (LNP) complexes show that the mRNA actively organises the internal lipid phase in a manner that correlates with transfection efficiency.^14–17^ Lipid encapsulation has likewise been applied to structured nucleic-acid assemblies, including DNA origami,^18^ and lipid-coated or lipid-associated nucleic acid nanostructures,^19,20^ indicating that defined nucleic-acid architectures are compatible with the lipid formulations used for mRNA delivery. Fixing the three-dimensional structure of the mRNA itself before use would remove an uncontrolled variable affecting both translation and delivery, and is the approach this work pursues.

DNA origami exploits the programmable base-pairing of nucleic acids to fold a long single-stranded scaffold into precise, nanometre-scale three-dimensional shapes defined entirely by the sequences of short complementary staple strands. Since the demonstration that DNA bundles can be programmed to twist and curve with controlled handedness,^24,25^ the field has developed a detailed understanding of how staple design determines the rigidity and solution shape of the folded object, supported by direct mechanical measurements showing that multi-helix bundles are substantially stiffer than duplex DNA and by finite-element tools that predict three-dimensional shape and flexibility directly from the design file. This combination of geometric programmability, mechanical rigidity, and predictable solution behaviour, together with the demonstrated capacity to decorate origami surfaces with proteins and other functional moieties at defined positions, has made structured nucleic-acid assemblies a versatile platform for biophysics and nanomedicine.^31,32^ The principle extends naturally from DNA to RNA: an RNA strand can itself serve as the scaffold, and single-stranded RNA origami can be folded cotranscriptionally as the molecule is synthesised, enabling genetic encoding and even intracellular expression of the resulting nanostructures. Recent advances have scaled cotranscriptional RNA origami to kilobase nanoscaffolds and produced dedicated design tools for RNA architectures, emphasising that the structural logic of origami is not confined to DNA.^36,37^ The scaffold need not be RNA alone: hybrid RNA–DNA origami, in which an RNA scaffold is folded by DNA staples, combine the addressability of DNA nanotechnology with an RNA cargo,^38^ including a functional messenger RNA scaffold.^39^ When the RNA scaffold is itself a protein-encoding messenger RNA, however, a distinct functional requirement emerges that has no analogue in structural DNA or RNA origami: the folded object must not only be geometrically and mechanically well-defined but must remain translationally competent.

We use DNA rather than RNA staples for three reasons: short ssDNA oligonucleotides are inexpensive and chemically robust, so a large staple set can be synthesised and stored without the handling constraints of RNA; DNA:RNA heteroduplexes form predictably with an mRNA scaffold of arbitrary sequence, whereas RNA nanostructure architectures are generally designed around defined tertiary motifs rather than applied to a pre-existing coding sequence. Furthermore, DNA staples are not substrates for the single-strand-specific ribonucleases from which the architecture is intended to protect the scaffold.

The fundamental conflict in this design is that the same staple hybridization that defines the structure and protects the backbone also occludes the transcript from the ribosome. Cap-dependent translation initiation in eukaryotes proceeds through a scanning mechanism in which the 43S preinitiation complex loads at the 5′ end of the mRNA and moves directionally along the leader sequence until it encounters the start codon.^43^ This process requires single-stranded RNA, since structured 5′ regions are known to impair initiation in proportion to their thermodynamic stability and proximity to the cap, because they impose barriers that the helicase eIF4A must resolve before the start codon becomes accessible.^44^ Toeprinting and ribosome profiling studies indicate that the 43S complex requires approximately 25–35 nucleotides (nt) of accessible RNA at the 5′ end for efficient loading. Below this threshold, initiation is severely compromised regardless of downstream sequence.^45^ An mRNA scaffold that is fully stapled along its entire length therefore satisfies the structural design constraint while violating the translational one. Leaving a defined region unstapled is the necessary compromise, but the design principles governing how long that overhang should be, what structural properties it must have, and how uridine chemistry modulates its behaviour are unknown.

This is the gap the present study addresses. Using a six-helix bundle (6HB) mRNA–DNA origami in which an EGFP-encoding mRNA serves as the scaffold, we first establish that the position of the unhybridized region determines translational output: exposing the proximal coding region raises output, whereas exposing the distal coding region alone does not. We then ask whether output increases with the length of the 5′ coding overhang across a 35–100 nt series, and find that it does not. Above the ribosomal initiation footprint, output declines with increasing overhang length, and no longer overhang exceeds the 35 nt reference. Finite-element modelling and imaging indicate that the constructs fold into 6HBs of comparable global morphology, and partition-function analysis identifies the predicted thermodynamic stability of the overhang as its closest computational correlate of output. That association cannot be separated from length, since longer overhangs are systematically more stable, and it is not robust to removal of either endpoint of the series; we therefore report it as a constraint on interpretation rather than as a mechanism. Finally, we show that a 35 nt overhang outperforms both the fully ORF-hybridized construct and a 100 nt overhang in unmodified, 5moU-and N1MeΨ-containing scaffolds, and that at a fixed 35 nt overhang N1MeΨ gives the highest output, followed by 5moU and then unmodified uridine. Together these observations identify coding-overhang accessibility and uridine chemistry as separable design parameters for translationally active mRNA–DNA hybrid origami, within the single open reading frame and single geometry examined here.

## Results

### Exposure of the proximal coding region increases translational output

To determine how the position of the unhybridized region within the coding sequence affects translational output, we designed a six-helix bundle (6HB) mRNA–DNA origami in which an EGFP-encoding mRNA serves as the scaffold strand, folded by ssDNA staple oligonucleotides (Figure 1A). The design is based on a previously reported architecture of Seitz et al. ^41^ modified to a 44 bp crossover spacing to accommodate the A-form geometry of the RNA:DNA heteroduplexes formed between scaffold and staples. The 35 nt window was chosen after an initial comparison in which exposing only the start codon gave lower output than exposing the first 35 coding nt (Supplementary Figure S1), consistent with the 25–35 nt of accessible RNA required for efficient 43S loading We prepared four constructs, each leaving a defined region of 35 nt of the coding sequence unhybridized: the coding sequence fully hybridized (0 nt coding overhang), the 3′ coding region unhybridized, the 5′ coding region unhybridized (reference), or both regions unhybridized simultaneously (Figure 1B). We included the double-unhybridized condition because we had previously found that both untranslated regions must remain accessible for efficient translation in mRNA–DNA origami ^41^ and because the closed-loop model of eukaryotic translation posits that eIF4G and poly(A)-binding protein bridge the 5′ cap and 3′ poly(A) tail into proximity to stimulate initiation. Staple hybridization at either end might therefore limit output independently. We note that this comparison does not test closed-loop formation directly, since the 3′ UTR and poly(A) tail remain unhybridized in all four constructs; it tests only whether exposure of the final 35 coding nucleotides contributes.

**Figure 1.**
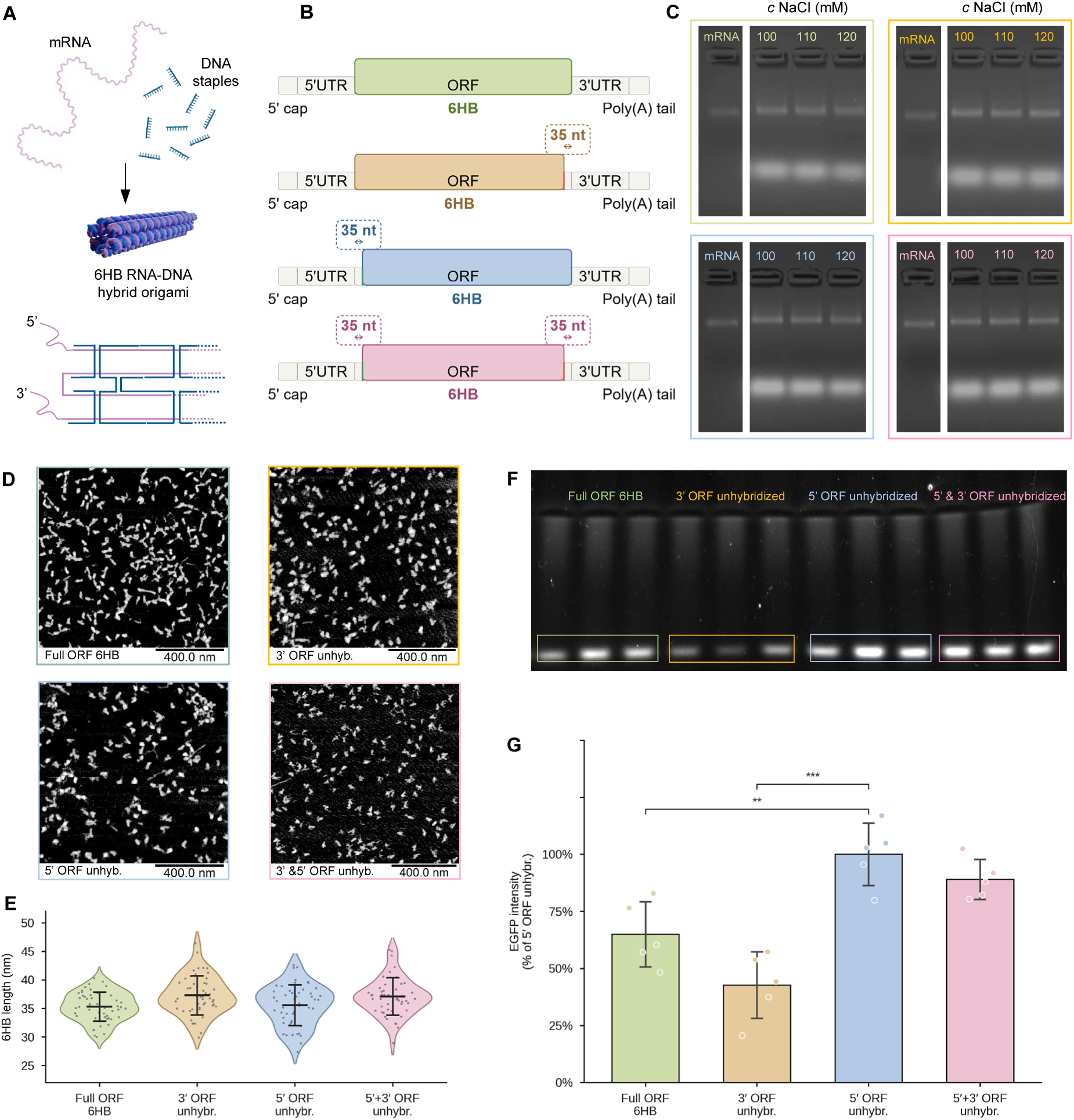
Exposure of the proximal coding region, not the distal, increases translational output in the hybrid six-helix bundle. (**A**) Design of the hybrid mRNA–DNA origami: an EGFP-encoding mRNA scaffold (top) is folded by short ssDNA staples into a six-helix bundle (6HB), shown as a three-dimensional model and as a crossover routing diagram. (**B**) Schematics of the four constructs, each leaving a defined 35 nt unhybridized region of the coding sequence (the coding overhang, i.e. the single-stranded portion of the scaffold left unhybridized by the staples, where the ribosome can access the transcript): fully ORF-hybridized (0 nt coding overhang), 3′ unhybridized, 5′ unhybridized (reference), and 5′ and 3′ unhybridized. In all constructs the 5′ UTR, 3′ UTR and poly(A) tail remain unhybridized. (**C**) Agarose gel electrophoresis of the four constructs folded at NaCl concentrations of 100, 110, and 120 mM. Folded species migrate as a compact, lower-mobility band relative to the naked mRNA scaffold (mRNA). (**D**) Representative atomic force microscopy (AFM) images of the folded constructs. (**E**) Violin plots of 6HB particle length measured by AFM, showing mean lengths of 35–37 nm with no significant differences between conditions. **(F**) Native PAGE of the translation reaction: in-gel Alexa 488 fluorescence reports the folded EGFP product. (**G**) EGFP fluorescence in rabbit reticulocyte lysate, normalised to the 5′-unhybridized reference (= 100%), with unfolded mRNA translated in parallel as a reference for the cost of folding; one-way ANOVA F(3,16) = 19.24, p < 0.0001, n = 5 independent folding and translation reactions. Bars show mean ± SD with individual replicates overlaid. **p < 0.01, ***p < 0.001 versus reference; all pairwise comparisons are given in Supplementary Table S2.

Agarose gel electrophoresis indicated successful folding of all four constructs, with folded species migrating as a compact, lower mobility band relative to the naked mRNA scaffold (Figure 1C). Atomic force microscopy revealed rod-shaped particles (Figure 1D) with mean lengths of 35–37 nm across all variants and no significant differences between conditions (Figure 1E), slightly exceeding the ∼30–34 nm predicted from the 6HB design. This modest overestimation is expected for AFM in air: the finite tip radius broadens and elongates features along the scan direction, the unhybridized single-stranded regions and short scaffold loops at the bundle ends extend beyond the rigid duplex body that the design length describes, and adsorption and partial flattening on the mica surface can increase the apparent length relative to the solution structure. The measured values are therefore consistent with the designed 6HB dimensions. An RNase A protection assay confirmed that staple hybridization shields the mRNA backbone from single-strand-specific nuclease digestion (Supplementary Figure S2).

We assessed translational output by cell-free translation in rabbit reticulocyte lysate (RRL), resolving the EGFP product by native PAGE. In-gel Alexa 488 fluorescence confirmed synthesis of folded EGFP across the constructs (Figure 1F). Quantified EGFP fluorescence (Figure 1G), normalized to the 5′ unhybridized reference construct (= 100%), showed that the position of the unhybridized region had a large and significant effect on translational output (p < 0.0001). The fully ORF-hybridized construct produced 65.0 ± 14.3% of reference output (p = 0.0031), confirming that hybridization of the coding sequence suppresses translation without abolishing it. Freeing the 3′ coding region alone did not recover output; it instead fell to 42.7 ± 14.6% (p < 0.0001 versus reference), the lowest value across all conditions. Freeing both the 5′ and 3′ regions simultaneously (89.0 ± 8.8%) was statistically indistinguishable from the 5′ unhybridized construct alone (p = 0.558).

Folding the transcript into a 6HB carries a translational cost that no arrangement of the unhybridized region recovers. An equimolar amount of unhybridized mRNA translated in parallel produced more EGFP than any folded construct, (Supplementary Figure S3). This is the expected consequence of occluding a transcript that must be read, and it means that the comparisons reported throughout this study describe relative performance within the origami series rather than parity with linear mRNA. The 3′-unhybridized construct is numerically lower than the fully ORF-hybridized control, which would be counterintuitive if exposure of any coding region were uniformly beneficial. What the comparison does establish is that exposing the last 35 coding nucleotides confers no measurable benefit, whereas exposing the first 35 does, and that adding 3′ exposure to a construct that already has 5′ exposure adds nothing. Accessibility of the proximal coding region is therefore the dominant positional determinant of output in this architecture. It is not, however, a requirement for translation as such: with the entire ORF hybridized, roughly two-thirds of reference output survives.

### Translational output declines as the 5′ coding overhang is extended beyond the initiation footprint

Above the initiation-footprint threshold, the length of the unhybridized 5′ overhang is the most obvious candidate variable for translational output. A longer free overhang increases the physical separation between the folded 6HB body and the site of initiation and lowers the probability that a scanning complex encounters the staple boundary before reaching the start codon, both of which would predict output rising with length. The 5′ UTR is identical and unhybridized in all constructs, so a longer coding overhang does not provide greater access to the cap or leader themselves. The staple boundary lies downstream of the AUG in every construct, so the scanning 43S complex should reach the start codon before encountering it in all cases. What a longer overhang does provide is a larger unstructured region downstream of the start codon, and the position-dependent model of mRNA structure does not predict that this should raise output ^11^.

We designed a series of constructs with unhybridized 5′ coding overhangs of 35, 50, 60, 70, 80 and 100 nt (Figure 2A), taking 35 nt as the series floor. As the overhang lengthens, the hybridized portion of the bundle shortens correspondingly, so we characterised the folded structures before comparing translational outputs. Finite-element modelling using SNUPI (Figure 2B) was used to compare the mechanical behaviour of the hybridized bundle across the series. Mean root mean square fluctuation (RMSF) reports the average per-residue thermal fluctuation across the hybridized 6HB body and serves as an index of global rigidity; maximum RMSF reports the peak displacement of the single most mobile residue and captures localized flexibility that the mean may obscure. Mean RMSF ranged from 0.748 to 1.218 nm (Figure 2C) and maximum RMSF from 2.759 nm at 80 nt to 8.357 nm at 50 nt (Figure 2D). The 50 nt construct combines a modest mean RMSF with the highest maximum in the series, consistent with a locally underconstrained crossover geometry at one bundle edge, while the 80 nt construct shows the lowest values of both. These are properties of the hybridized body only; the model does not describe the single-stranded overhang, and we do not use them to interpret translational output.

**Figure 2.**
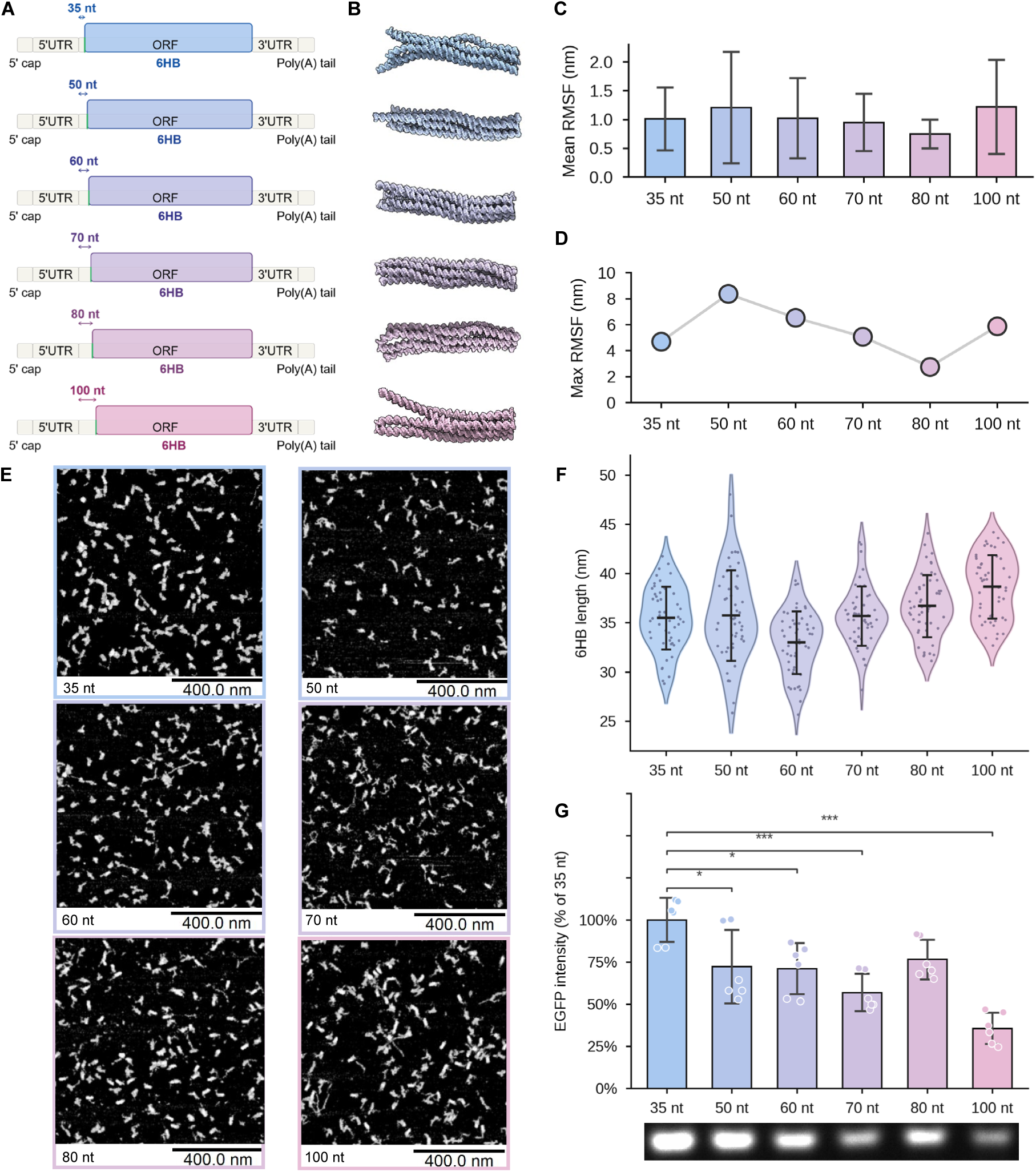
Translational output declines as the 5′ coding overhang is extended beyond the initiation footprint. (A) Schematics of the construct series, with unhybridized 5′ coding overhangs of 35, 50, 60, 70, 80 and 100 nt. (B) Three-dimensional models of each construct from finite-element modelling (SNUPI). (C) Mean root mean square fluctuation (RMSF) of the hybridized 6HB body across the series, reporting global mechanical rigidity. (D) Maximum RMSF across the series, reporting the peak local fluctuation. RMSF values in (C) and (D) describe the hybridized body only; the model does not describe the single-stranded overhang, and these values are reported as structural characterisation rather than as an explanation of translational output. (E) Representative AFM images of each construct (scale bars, 400 nm). (F) Violin plots of 6HB particle length by AFM, confirming mean lengths of 35–38 nm with no significant differences between conditions. (G) EGFP fluorescence in rabbit reticulocyte lysate, normalised to the 35 nt construct (= 100%), with the corresponding native PAGE below; one-way ANOVA F(5,30) = 13.51, p < 0.0001, n = 6 independent folding and translation reactions. Bars show mean ± SD with individual replicates overlaid. Asterisks denote comparisons against the 35 nt reference only (*p < 0.05, ***p < 0.001); the complete 15-pair comparison matrix is given in Supplementary Table S3c.

AFM imaging (Figure 2E) confirmed rod-shaped particles across the series, with mean lengths of 35–38 nm and no significant differences between conditions (Figure 2F). The particles appeared somewhat less sharply defined than a typical all-DNA 6HB bundle, consistent with the architecture rather than with folding defects: the untranslated regions and the 5′ coding overhang are single-stranded by design and contribute local conformational disorder at the scaffold termini, and the hybridized core is formed by A-form RNA:DNA heteroduplexes rather than B-form DNA. Together with the folding gels (Supplementary Figure S4), these data indicate that all constructs fold into well-defined 6HB nanoparticles of comparable global morphology and apparent length.

The translational data show no monotonic increase (Figure 2G, p < 0.0001). Output, normalized to the 35 nt reference (= 100%), declined as the overhang lengthened: 72.3 ± 21.8% at 50 nt (p = 0.023), 71.2 ± 15.1% at 60 nt (p = 0.017), 57.0 ± 11.1% at 70 nt (p = 0.0002), 76.5 ± 11.8% at 80 nt (p = 0.077) and 35.7 ± 9.3% at 100 nt (p < 0.0001) (Supplementary Tables S3a and S3b). No construct exceeded the 35 nt reference, and the 100 nt construct produced the lowest output.

The response is not a graded decline. Output within the 50–80 nt interval does not depend on overhang length: the four constructs in this range are statistically indistinguishable from one another (p = 0.179), and a test for monotone trend restricted to them is likewise non-significant (p = 0.959; Supplementary Table S3d). Treating this interval as a single regime, the series resolves into three levels: 100% at 35 nt, 69.2% across the pooled 50–80 nt interior, and 35.7% at 100 nt, with all three differing significantly from one another (p ≤ 0.0002 for every pairwise comparison; Supplementary Table S3d). This description fits the construct means considerably better than a continuous decline in length (R² = 0.905 versus 0.737). The numerically higher value at 80 nt falls within the interior regime and is not distinguishable from the 50, 60 or 70 nt constructs (p = 0.995, 0.986 and 0.196 respectively; Supplementary Table S3c).

Output therefore falls in two discrete steps rather than in proportion to overhang length. Exposing coding sequence beyond the ribosomal initiation footprint incurs a penalty that is already complete at 50 nt and does not increase across the 50–80 nt range, followed by a second, larger penalty at 100 nt. Across the full 0–100 nt range, combining the fully ORF-hybridized construct (65.0% of the 35 nt condition) with this series, output is highest at 35 nt and lower at both extremes. This shape is not unexpected: low structure across the leader and the first codons favours initiation, while greater structure further into the coding sequence is not detrimental and can extend the period over which a transcript is translated.¹¹ The observation of interest is therefore not that output fails to rise with overhang length, which the position-dependent model never predicted, but that the cost of over-exposure is discontinuous.

### Predicted overhang folding stability does not provide a determinant independent of length

To test whether sequence-level structural properties of the free overhang could account for the length dependence, we computed partition-function statistics for each unmodified-uridine overhang sequence using RNAfold (ViennaRNA 2.7.2).^46^ This model provides the minimum free energy (ΔG), the frequency of the minimum-free-energy (MFE) structure in the Boltzmann ensemble, and the ensemble diversity (Figure 3A, Supplementary Table S4; predicted MFE structures in Supplementary Table S6).

**Figure 3.**
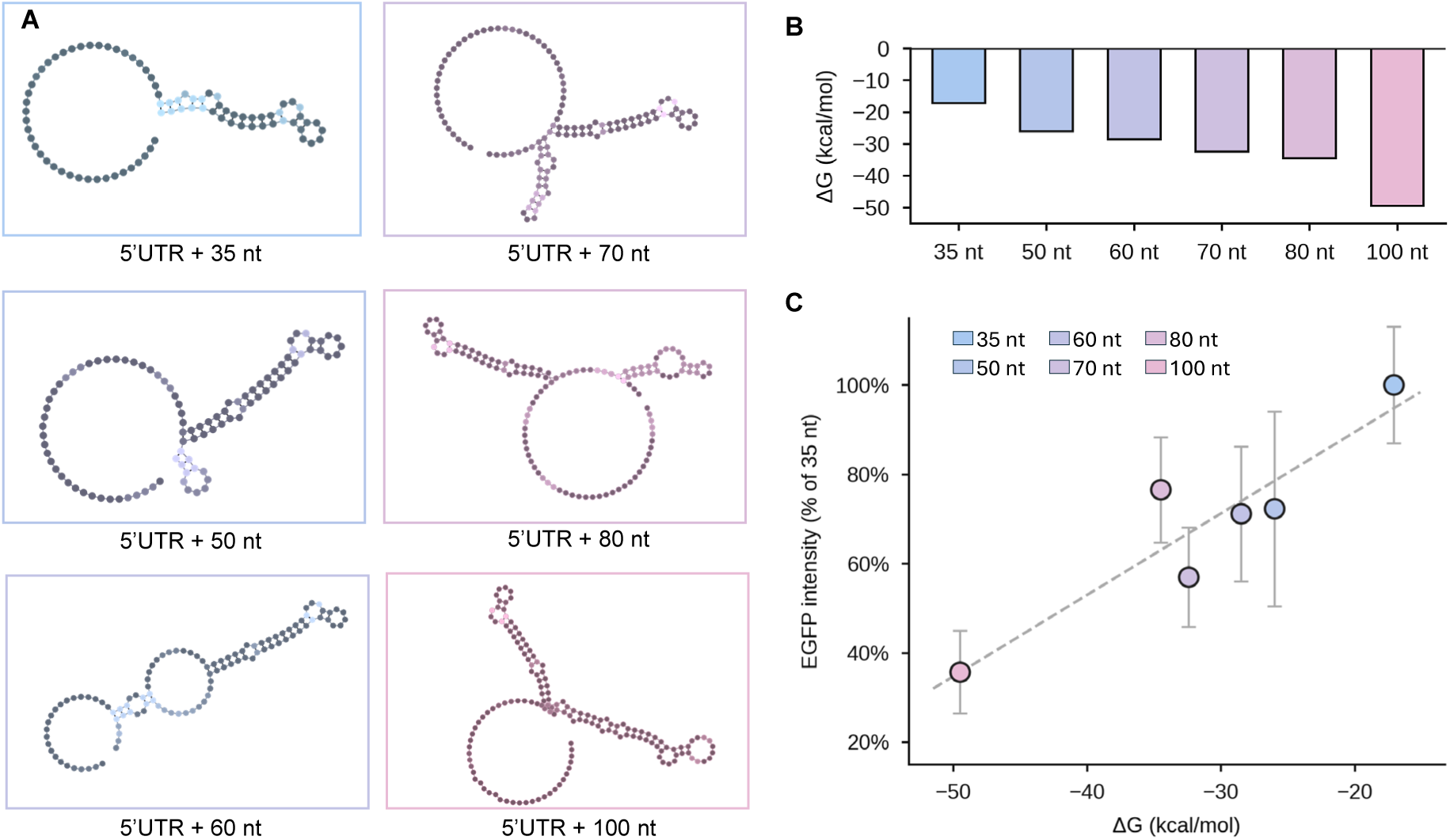
Predicted overhang folding stability co-varies with overhang length and does not provide an independent determinant of output. (A) Minimum-free-energy (MFE) secondary structures of the unmodified-uridine overhang sequences (5′ UTR + 35, 50, 60, 70, 80 or 100 nt of coding sequence), predicted by RNAfold (ViennaRNA 2.7.2). (B) Predicted folding free energy (ΔG) across the length series. (C) Translational output (EGFP, % of 35 nt) versus ΔG for the six constructs; Pearson r = +0.920, R² = 0.846, p = 0.009. The dashed line is a linear fit. Overhang length and ΔG co-vary almost perfectly across the series (r = −0.981), so this correlation cannot be interpreted independently of length; it is also leverage-dependent, falling to r = +0.798 (p = 0.105) when the 100 nt construct is omitted (Supplementary Table S5b)

Thermodynamic stability increased monotonically with overhang length, ΔG ranging from −17.10 kcal/mol at 35 nt to −49.50 kcal/mol at 100 nt (Figure 3B). Across the six constructs, ΔG was the strongest correlate of translational output (Pearson r = +0.920, R² = 0.846, p = 0.009; Figure 3C, Supplementary Table S5), with less stable overhangs associating with higher output, whereas MFE frequency (r = +0.356, p = 0.488) and ensemble diversity (r = +0.258, p = 0.621) showed no association.

This correlation cannot be read as an independent structural effect, for two reasons. First, overhang length and ΔG co-vary almost perfectly across the series (r = −0.981, p = 0.0005), because longer overhangs are systematically more stable, and length itself correlates with output (r = −0.859, p = 0.028); the two variables cannot be separated within this construct set. Second, the fit is leverage-dependent. Leave-one-out analysis (Supplementary Table S5b) shows that omitting the 100 nt construct, which lies far from the remaining five in both ΔG and output, reduces the correlation to r = +0.798 (p = 0.105), and among the 35–80 nt constructs alone the relationship does not reach significance. The correlation is therefore consistent with a model in which a more stable overhang structure impedes translation, but it neither establishes overhang stability as a determinant independent of length nor rests on more than the endpoint of the series.

We also examined whether the translational differences instead arise from accessibility of the translation initiation region specifically, computing the predicted unpaired probability of the start codon and first five codons from the full-construct partition function (Supplementary Table S4). This accessibility was essentially invariant across the 50–100 nt constructs (0.286– 0.290) despite a more than two-fold range in translational output among them, and its apparent correlation with output across the full series (r = +0.792) does not reach significance (p = 0.060) and reflects only the shorter 35 nt construct. The invariance is expected, since the ribosome loading zone is sequence-identical across all constructs and they differ only in the length of downstream coding sequence left unhybridized. We note that this analysis relies on the same predictive tool whose limitations we discuss below, and it is therefore reported as consistent with, rather than as demonstrating, the absence of an initiation-region effect.

Finally, we examined the full Boltzmann structural ensemble of each overhang by stochastic sampling. The number of independent structural domains was a robust ensemble-level feature, but it scaled with overhang length and did not separate output from length (Supplementary Tables S7 and S8): constructs with near-identical domain distributions translated very differently, and none of the domain metrics correlated with output beyond what length already accounts for.

The computational analysis therefore identifies no structural property of the overhang that accounts for translational output independently of its length. We report this as a negative result. Within this construct set, overhang length and predicted folding stability are experimentally inseparable, and no finer ensemble feature we examined separates them. Establishing whether overhang structure has any effect distinct from length will require constructs matched in length but differing in predicted stability, or direct experimental probing of overhang structure within the assembled origami.

### Uridine chemistry sets the magnitude of translational output

Having characterised the overhang length response for the unmodified scaffold, we asked next whether uridine chemistry alters this behaviour. We first tested two clinically relevant modifications, 5moU and N1MeΨ, using the fully ORF-hybridized (0 nt), 35 nt and 100 nt constructs (Figure 4A, 4C), with output normalised to each modification’s own 35 nt condition (= 100%; n = 6 per condition). Both modified scaffolds folded into intact 6HB structures, observed by agarose gel electrophoresis (Supplementary Figure S5) and, for the 35 nt construct of each modification, by AFM (Supplementary Figure S6). Both 5moU and N1MeΨ alter the base-pairing and stacking thermodynamics of the scaffold, which could reduce folding efficiency for a staple set designed against unmodified RNA. Folding efficiency was nonetheless comparable to the unmodified constructs across all conditions tested. A single staple design can therefore be applied across all three chemistries without re-optimisation.

**Figure 4.**
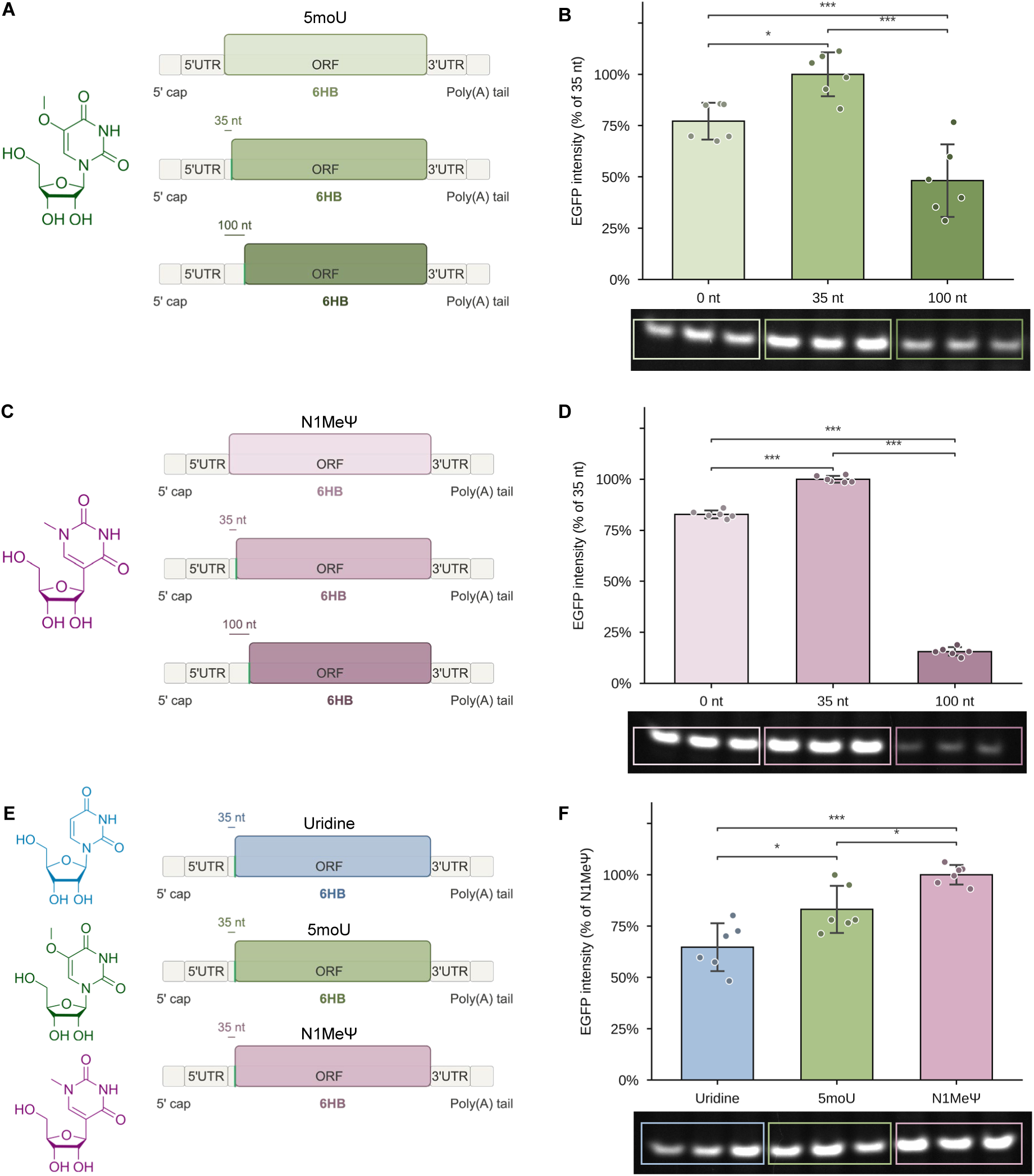
Uridine chemistry sets the magnitude of translational output. (A) Schematics of the 5-methoxyuridine (5moU) constructs at 0, 35 and 100 nt overhang length. (B) EGFP fluorescence for the 5moU constructs with corresponding native PAGE, normalised to the 5moU 35 nt construct (= 100%). (C) Schematics of the N1-methylpseudouridine (N1MeΨ) constructs at 0, 35 and 100 nt. (D) EGFP fluorescence for the N1MeΨ constructs with native PAGE, normalised to the N1MeΨ 35 nt construct (= 100%). (E) Schematics of the cross-modification comparison at a fixed 35 nt overhang for unmodified uridine, 5moU and N1MeΨ. (F) EGFP fluorescence for the cross-modification comparison with native PAGE, normalised to N1MeΨ (= 100%); one-way ANOVA F(2,15) = 19.29, p < 0.0001. In (B), (D) and (F) unfolded mRNA of the corresponding chemistry was translated in parallel as a reference for the cost of folding. All panels: n = 6 independent folding and translation reactions; bars show mean ± SD with individual replicates overlaid; *p < 0.05, **p < 0.01, ***p < 0.001 (Tukey HSD).

Under 5moU (Figure 4B), output depended strongly on the coding overhang (p < 0.0001). The 35 nt construct gave the highest output, exceeding both the fully ORF-hybridized construct, which reached 77.1 ± 9.0% of that value (p = 0.021), and the 100 nt construct, which reached 48.2 ± 17.7% (p < 0.0001). The 100 nt construct also fell below the fully ORF-hybridized construct (p = 0.0042). Under N1MeΨ (Figure 4D) the ordering was the same but the separation greater: 82.8 ± 1.9% for the fully ORF-hybridized construct and 15.6 ± 2.1% for the 100 nt overhang, with all three comparisons significant at p < 0.0001.

The direction of the response is common to both modifications, but its magnitude is not. Analysing the two modified series together as a two-way design, chemistry and overhang length both have strong main effects, and their interaction is significant (p < 0.0001; Supplementary Table S11c). Decomposing that interaction locates it entirely in the over-exposed condition. The benefit of exposing the proximal coding region is quantitatively indistinguishable between chemistries (p = 0.299), whereas the penalty for exposing 100 nt is not (p < 0.0001): output falls to 45.8% of the 35 nt reference under 5moU but to 15.5% under N1MeΨ. This is not an artefact of normalising to each chemistry’s own 35 nt construct, since the same difference is obtained when the 100 nt construct is compared directly with the fully ORF-hybridized construct, a contrast that does not involve the 35 nt reference (p < 0.0001). Sensitivity to an over-long overhang is therefore roughly threefold greater for N1MeΨ. Because both modifications alter base-pairing and stacking thermodynamics, a plausible reading is that the same exposed sequence folds more stably in a modified scaffold, and that the penalty for over-exposure scales with that stability.^47,48^ This is consistent with the association between predicted overhang stability and output reported above, and it arises here from a comparison in which overhang length is held constant.

To compare uridine chemistries on a common reference, we then measured translational output at a fixed 35 nt overhang across all three scaffolds, where overhang length and sequence are identical and only the scaffold modification differs (Figure 4E). Normalised to the highest-performing chemistry (N1MeΨ = 100%), unmodified uridine translated at 64.7% and 5moU at 83.1% (Figure 4F; p < 0.0001). Both modifications significantly increased output relative to the unmodified scaffold (5moU, p = 0.014; N1MeΨ, p < 0.0001), and N1MeΨ significantly exceeded 5moU (p = 0.025). The ranking is consistent with the established enhancement of translational output by nucleoside modification in linear mRNA^4,8^, and demonstrates that this enhancement is retained within the hybrid origami architecture. Uridine chemistry therefore sets the magnitude of output, while the response to overhang position and length is common to all three scaffolds. Raw replicate values for all conditions are provided in Supplementary Tables S9–S11.

## Discussion

The design of a translationally competent mRNA–DNA origami is governed by a conflict between two requirements: staple hybridization defines the structure and shields the backbone, while the ribosome requires single-stranded RNA. The experiments reported here resolve part of that conflict into a positional rule. Accessibility of the proximal coding region determines output, whereas accessibility of the distal coding region does not, and once the proximal region is exposed, additional exposure at the 3′ end contributes nothing further. This is consistent with cap-dependent scanning being the rate-limiting step in this architecture: the 43S preinitiation complex loads at the 5′ end and translocates directionally, and it is the region it must traverse before reaching the start codon that determines whether initiation succeeds. ^43^ Because the 3′ UTR and poly(A) tail remain single-stranded in every construct, these data do not address closed-loop formation; what they establish is that hybridizing the final 35 coding nucleotides carries no measurable penalty, which is useful design latitude.

The most informative number in the study may be the easiest to overlook. With the entire open reading frame hybridized, the construct still translates at 65% of the 35 nt reference under unmodified uridine, and at 77% and 83% under 5moU and N1MeΨ respectively. Proximal exposure is therefore beneficial rather than required, and folding the transcript into a six-helix bundle does not render it inert. Substantial output from a fully ORF-hybridized construct implies that elongating ribosomes displace staples as they translate, since the ribosomal mRNA entry channel accommodates only single-stranded RNA. The folded state is therefore likely to be transient under translating conditions. This describes how the architecture works rather than a limitation of it. The structure defines the object during handling, formulation and delivery, and is resolved by the ribosome at the point of use. We did not assess construct integrity after translation, and the folded state should not be assumed to survive it.

Above the initiation footprint, output declines as the exposed coding region lengthens. This is not the counterintuitive result it may appear. The 5′ UTR is unhybridized in every construct, so a longer coding overhang confers no additional access to the cap or leader, and the staple boundary lies downstream of the start codon throughout; what a longer overhang provides is a larger unstructured region within the coding sequence, which the position-dependent model of mRNA structure does not predict should raise output. ^11^ Within the 35–100 nt series the fall is not gradual but stepwise: output is indistinguishable across 50–80 nt before dropping again at 100 nt, so the penalty for exceeding the initiation footprint is a threshold rather than a gradient.

Predicted thermodynamic stability of the overhang is its closest computational correlate, and less stable overhangs translate better, which would be consistent with eIF4A resolving 5′ structure during scanning. ^44^ But stability and length co-vary almost perfectly across a series in which longer overhangs are necessarily more stable, and the correlation is not robust to removal of either endpoint. Neither ensemble diversity, minimum-free-energy frequency nor the distribution of independent folded domains separates output from length. Therefore it is not possible to assign a structural mechanism to the decline. A more fundamental obstacle limits what prediction can contribute here: RNAfold computes the structure of a free RNA fragment, whereas in the assembled origami the overhang is anchored at a staple junction against a rigid A-form helix, a constraint that partition-function calculations do not represent and that may alter base-pairing at the UTR–ORF boundary. Separating length from structure will require constructs matched in length but differing in predicted stability, direct chemical probing of the overhang within the assembled object, and folding tools that account for junction context.

Uridine chemistry acts on the magnitude of output rather than on the response to overhang geometry. At a fixed 35 nt overhang, N1MeΨ outperforms 5moU, which outperforms unmodified uridine, reproducing within a folded nanostructure the ranking established for linear transcripts. ^49^ Because these measurements are cell-free, the enhancement cannot derive from evasion of innate immune sensing. It is consistent with two mechanisms characterised in lysate: suppression of eIF2α phosphorylation, which unmodified mRNA induces and modified mRNA avoids, ^50^ and an eIF2α-independent increase in ribosome density and loading attributed to altered interactions at the decoding centre. ^10^ That the same 0–35–100 nt response shape holds across all three chemistries, while their absolute outputs differ, indicates that overhang geometry and nucleoside chemistry are separable and potentially additive design parameters. A single staple set folds all three scaffolds without re-optimisation, so a modified transcript can be substituted into an existing design without redesigning the structure.

Every folded construct translates less efficiently than the same mRNA left unhybridized. This is the expected cost of occluding a transcript that must be read, and whether it is acceptable depends on a comparison that endpoint measurement in lysate cannot make. A 90-minute reaction in reticulocyte lysate weights initiation efficiency heavily and transcript survival minimally, since the system imposes no meaningful nuclease pressure and no delivery step. In a cellular context the balance may differ: hybridization demonstrably shields the backbone from single-strand-specific nuclease attack, and a construct that initiates less efficiently but persists longer can produce more total protein than one translated efficiently and then degraded. This is precisely the argument the position-dependent model makes for downstream secondary structure, where increased structure lowers initiation rate while extending functional half-life and raising cumulative yield. ^11^ Whether hybrid origami sits on the favourable side of that trade-off is the question that determines the utility of the architecture, and it cannot be settled by the measurements reported here.

Answering it is presently limited by delivery rather than by design. Standard cationic-lipid transfection reagents are formulated and validated for linear single-stranded mRNA, and no standardized reagent exists for delivering a rigid nanostructure of this size intact. Complexation conditions optimised for a flexible polyanion are different for a folded six-helix bundle. Lipid encapsulation of structured nucleic-acid assemblies is achievable but requires bespoke formulation. ^18–20^ There is also a second, interpretive difficulty: constructs in this series differ in staple content, hybridized length and exposed single-stranded RNA, so uptake, endosomal escape and intracellular stability would be expected to differ between them. Without independently matched delivery, differences in cellular output could not be attributed to coding-overhang accessibility. Formulation development for this construct class is therefore a prerequisite for the relevant cellular experiments rather than a downstream application of them.

## Conclusions

We have defined how the placement and extent of unhybridized coding sequence govern translation from an mRNA–DNA hybrid 6HB. Exposure of the proximal coding region determines output; exposure of the distal region does not, and extending the exposed region beyond the ribosomal initiation footprint progressively reduces translation rather than improving it. A 35 nt proximal overhang is the best-performing configuration among those tested, and it holds across unmodified, 5moU-and N1MeΨ-containing scaffolds folded with a single staple set. Uridine chemistry sets the magnitude of output, with N1MeΨ highest, without altering the response to overhang geometry, so the two parameters can be optimised independently.

For design, the operative rule is to expose the initiation footprint and no more, and to select uridine chemistry separately. For the field, the question that follows is whether the initiation penalty measured in a cell-free system is offset by extended transcript survival in a cellular one. Both require a delivery route that preserves the folded structure, making formulation development for hybrid mRNA–DNA origami the immediate priority.

## Methods

### Design of the mRNA–DNA hybrid origami

The hybrid origami structures were designed on a honeycomb lattice in caDNAno ^51^ and simulated with CanDo ^27^ (27, Kim et al. 2012). The fully ORF-hybridized origami was based on the six-helix bundle (6HB) D2 architecture reported by Seitz et al.^41^, with minor modifications. The crossover spacing was adjusted from the original 42 bp to 44 bp, corresponding to a change from 10.5 to 11 bp per helical turn to match the A-form geometry of the RNA:DNA heteroduplexes formed between the mRNA scaffold and the ssDNA staples. A-form helices are shorter and wider than B-form, with a helical rise of approximately 2.55 Å per base pair, a diameter of roughly 2.3 nm and about 11 base pairs per turn rather than the ∼10.5 of B-form DNA. Placing crossovers at an integer multiple of the A-form helical repeat was intended to position staple crossovers on the correct helical face to connect adjacent helices without introducing torsional strain.

The fully ORF-hybridized design was then modified by leaving a defined region of the open reading frame (ORF) unhybridized at the 5′ end, the 3′ end, or both, with varying numbers of nucleotides. The ORF is defined here as the coding sequence from the start codon to the stop codon. The 5′ and 3′ untranslated regions (UTRs) of the scaffold, the poly(A) tail, and short scaffold loops (7–8 nt) at both ends of each structure were left unhybridized in all constructs; “fully ORF-hybridized” therefore refers to the coding sequence only. Each construct was designed independently in caDNAno rather than by omitting staples from a single full-length design. For each variant, the 5′ boundary of the hybridized region was moved downstream by the required number of nucleotides and the scaffold routing re-generated, so that the shortened 6HB body retains a complete and regular crossover pattern up to the new boundary. The staple strands for each variant are listed in **Supplementary Table S1**, while complete caDNAno design files for all constructs are available from the authors on request.

A 997 nt EGFP-encoding mRNA (TriLink Biotechnologies), comprising a 720 nt ORF together with the 5′ cap, UTRs, and poly(A) tail, was used as the scaffold. The scaffold is a proprietary commercial construct and sequence details are available from the manufacturer. Staple strands were purchased from Integrated DNA Technologies.

### Folding and assembly

The folding protocol was adapted from Wang et al.^42^ Folding reactions contained 50 nM mRNA scaffold and a 10-fold molar excess of staple strands (500 nM each) in 1× folding buffer (10 mM Tris), prepared with nuclease-free water (Thermo Fisher Scientific). To identify optimal folding conditions, the NaCl concentration was screened across seven values (70–130 mM) for all constructs and both annealing protocols (Supplementary Figure S7).

Folding was carried out in a thermal cycler (Applied Biosystems ProFlex PCR System) using one of two annealing protocols adapted from Wang et al., in which the optimal maximum folding temperature was found to lie between 50 and 60 °C: either isothermal incubation at 55 °C for 30 min, or a linear temperature ramp from 50 °C to 20 °C at 0.125 °C min⁻¹ over 4 h. After folding, structures were either incubated on ice for 15 min or stored immediately at 4 °C. All structures were used freshly prepared or at most one day after folding to avoid storage-related instability. Constructs used in all subsequent experiments were folded at 110 mM NaCl in 10 mM Tris using the 50 °C to 20 °C ramp.

### Atomic force microscopy

Atomic force microscopy (AFM) was used to characterize folding, with a protocol adapted from Seitz et al. Samples were prepared at 7.5 nM unpurified origami in 16 mM MgCl₂ (50 µl total volume). A 20 µl aliquot was deposited onto freshly cleaved mica and incubated for 5 min to allow attachment of the negatively charged structures to the mica surface via Mg²⁺ bridging. The surface was then washed three times with nuclease-free water to remove unbound material and dried under a stream of nitrogen. Samples were imaged in air on a Dimension Icon AFM (Bruker) in ScanAsyst mode using ScanAsyst Air cantilevers (Bruker). Images were processed in NanoScope Analysis v3.0, and particle lengths were measured in ImageJ.

### Agarose gel electrophoresis

Folding and structural integrity were assessed by agarose gel electrophoresis. Gels contained 3.5% (w/v) agarose in 1× TAE with 11 mM MgCl₂ and were stained with 0.46 µg ml⁻¹ ethidium bromide; the running buffer was 1× TAE with 11 mM MgCl₂. For folding analysis, samples were prepared at 12.5–25 nM and combined with 6× loading dye (40% sucrose) at a 1:5 volume ratio. For stability assays, samples were prepared at 7.5 nM (RNase A), and 40% sucrose without dye was added to prevent overlap of dye bands with digestion-assay bands. A 10 µl aliquot of each sample was loaded per lane. Gels were run at 90 V for 45 min in an ice bath and imaged on a ChemiDoc MP system (Bio-Rad) under UV illumination (ethidium bromide channel); band intensities were quantified in ImageJ.

### In vitro translation

Translational competence was assessed using a Rabbit Reticulocyte Lysate System (Promega) according to the manufacturer’s protocol with minor modifications. Each 50 µl reaction contained 17 µl rabbit reticulocyte lysate, 27 µl nuclease-free water, and 1 µl of an amino acid mixture prepared by combining equal volumes of a methionine-free and a leucine-free amino acid mixture (each 1 mM) to provide a complete set of amino acids. Folded origami or unfolded mRNA was added to a final concentration of 5 nM; unfolded mRNA was included in every experiment as a reference for the translational cost of folding. Reactions were incubated at 37 °C for 90 min and then placed on ice to terminate translation.

Translation products were resolved by native PAGE. Each reaction was combined 1:1 with 2× native PAGE loading dye (62.5 mM Tris, 40% v/v glycerol, 0.01% w/v bromophenol blue), and 10 µl was loaded onto a 4–20% Mini-PROTEAN TGX precast gel (Bio-Rad). Gels were run in 25 mM Tris, 192 mM glycine (pH 8.3) at 200 V for 45 min in an ice bath and imaged on a ChemiDoc MP system (Bio-Rad) using the Alexa 488 channel. The readout throughout is in-gel fluorescence of the folded, chromophore-matured EGFP product, and therefore reports the combined outcome of translation, co-translational folding and chromophore maturation rather than peptide synthesis alone. Band intensities were quantified in ImageJ. Each replicate reported in this study represents an independent folding reaction and an independent translation reaction; replicates are not repeated measurements or repeated quantifications of a single preparation.

### Computational RNA secondary structure analysis

Secondary structure prediction was performed with the ViennaRNA package^46^ (version 2.7.2) using the Turner 2004 nearest-neighbour parameters at 37 °C.^52^ Each construct sequence comprised the 48 nt 5′ UTR of the mRNA followed by the first 35, 50, 60, 70, 80, or 100 nt of the EGFP ORF, corresponding to the unhybridized overhang of each construct. For each sequence, the minimum free energy (ΔG) and minimum-free-energy structure were computed, and the partition function was evaluated to obtain the frequency of the minimum-free-energy structure within the Boltzmann ensemble and the ensemble diversity (mean base-pair distance between sampled structures).^53^ A large language model (Claude, Anthropic, accessed June 2026) was used to assist with the development of computational analysis code. All outputs were reviewed and validated by the authors.

Accessibility of the translation initiation region was assessed from the same full-construct partition function as the mean unpaired probability of the start codon and the first five codons of the ORF (positions 49–63). Because this region is identical in sequence across all constructs, it serves as a control for whether predicted initiation-region structure varies across the series.

To characterize the structural ensemble beyond the single minimum-free-energy structure, 5000 structures were drawn from the Boltzmann ensemble of each construct by stochastic backtracking with unique multiloop decomposition enabled. Each sampled structure was decomposed into independent base-paired domains, defined as maximal base-paired segments separated by unpaired nt, and the distribution of domain counts across the ensemble was tabulated.^54^

### Finite-element modelling

The mechanical properties of the folded constructs were predicted using the Structured NUcleic acids Programming Interface (SNUPI), a finite-element framework for nucleic acid nanostructures.^28^ Each origami was modelled in equilibrium to extract its structural properties, including the root mean square fluctuation (RMSF) of the hybridized regions, obtained by normal mode analysis. This model describes the double-stranded helical bundle formed by the hybridized staple regions; it does not describe the conformational dynamics of the single-stranded free overhang. The nonlinear mechanical behaviour of both single-stranded DNA and single-stranded RNA was included, and the base-pair and crossover step parameters were derived from all-atom molecular dynamics simulations as reported in the literature. Simulations used the standard SNUPI parameter set: base-pair and crossover steps with a coefficient function of order 2; for single-stranded DNA, a contour length per nt of 0.38 nm (short) and 0.68 nm (long), a persistence length of 0.67 nm, stretching rigidities of 710 pN (stretched) and 15 pN (relaxed), and a coefficient function of order 3; for single-stranded RNA, a contour length per nt of 0.383 nm (short) and 0.606 nm (long), a persistence length of 0.606 nm, and stretching rigidities of 1512 pN (stretched) and 8 pN (relaxed); and electrostatic interactions at 300 K and 300 mM NaCl, with an effective charge of 1 e per base pair, a cutoff distance of 2.5 nm, and a coefficient function of order 1.^42^

### Statistical analysis

Translational output was quantified from native PAGE band intensities and analysed by one-way analysis of variance (ANOVA) followed by Tukey’s honestly significant difference (HSD) test across all pairwise comparisons. Comparisons against a single reference condition are reported alongside the complete pairwise matrix rather than in place of it. Homogeneity of variance was assessed by Levene’s and Bartlett’s tests and normality by the Shapiro–Wilk test. Where a homogeneity or normality assumption was not fully satisfied, Games–Howell comparisons, which estimate the standard error separately for each pair, are additionally reported in the Supplementary Information as a sensitivity analysis; Tukey values are those reported in the text and figures. Correlations between computed sequence metrics and translational output were evaluated across the six constructs (n = 6) using Pearson correlation on construct-level mean values, with two-tailed p-values, and the sensitivity of each correlation to individual constructs was assessed by leave-one-out analysis. Data are reported as mean ± standard deviation, with significance denoted as *p < 0.05, *p < 0.01, p < 0.001. Analyses were performed in Python (SciPy 1.17, statsmodels 0.14).

## Acknowledgements

This work has received funding from the European Research Council (ERC) under the European Union’s Horizon 2020 research and innovation programme (Grant Agreement No. 101002258), Research Council of Finland Centers of Excellence Programme, Life Inspired Hybrid Materials (LIBER) project number 346110 and the Research Council of Finland Flagship Programme for the Gene, Cell and Nano Therapy Competence Cluster for the Treatment of Chronic Diseases (GeneCellNano). We acknowledge the provision of facilities and technical support by Aalto University Bioeconomy Facilities, OtaNanoNanomicroscopy Center (Aalto-NMC) and Micronova Nanofabrication Center.

## Author contributions

Carmine D’Amico (Conceptualization [supporting], Data curation [lead], Formal analysis [lead], Investigation [lead], Methodology [lead], Visualization [lead], Writing – original draft [lead], Writing – review & editing [equal]), Miska Mykkänen (Investigation [supporting], Formal analysis [supporting], Writing – review & editing [supporting]), Sharon Saarinen (Investigation [supporting], Formal analysis [supporting], Writing – review & editing [supporting]), Ville Säkkinen (Investigation [supporting]), and Mauri A. Kostiainen (Conceptualization [lead], Funding acquisition [lead], Project administration [lead], Resources [lead], Supervision [lead], Writing – review & editing [equal]).

## Data Availability

All data underlying this article are available in the article and its online Supplementary Information. This includes the raw translation, atomic force microscopy, and gel-quantification values, the predicted secondary-structure metrics, and the full staple oligonucleotide sequences.

The computational scripts used for the secondary-structure analysis are available at Zenodo (https://doi.org/10.5281/zenodo.20830430). Secondary-structure prediction used the freely available ViennaRNA package version 2.7.2 (https://www.tbi.univie.ac.at/RNA/). The EGFP-encoding scaffold mRNA (TriLink Biotechnologies) and staple oligonucleotides (Integrated DNA Technologies) are commercially available.

