## Supporting information for "Impact of Coding Region Accessibility and Uridine Chemistry on Translation in mRNA–DNA Hybrid Origami"

### Supplementary Table S1. Staple sequences

*DNA staple strands used to fold each mRNA–DNA hybrid origami construct.*

#### S1 summary. Staple composition of each construct

| Panel | Construct | Staples | Total nt | Length range (nt) | Mean length (nt) |
|---|---|---|---|---|---|
| S1a | Fully ORF-hybridized | 23 | 663 | 21–43 | 28.8 |
| S1b | 3′ unhybridized | 23 | 642 | 21–37 | 27.9 |
| S1c | 5′ unhybridized (35 nt overhang) | 21 | 623 | 21–43 | 29.7 |
| S1d | 5′ and 3′ unhybridized | 21 | 594 | 21–42 | 28.3 |
| S1e | 50 nt overhang | 22 | 634 | 21–43 | 28.8 |
| S1f | 60 nt overhang | 22 | 624 | 21–43 | 28.4 |
| S1g | 70 nt overhang | 22 | 614 | 18–43 | 27.9 |
| S1h | 80 nt overhang | 21 | 604 | 18–43 | 28.8 |
| S1i | 100 nt overhang | 20 | 583 | 21–50 | 29.1 |

#### S1a. Fully ORF-hybridized (0 nt coding overhang) — 23 staples

| # | Sequence (5′→3′) | nt | # | Sequence (5′→3′) | nt |
|---|---|---|---|---|---|
| 1 | TCGAACTTCACCTCGGCGCGGGTGGTGCAG | 30 | 13 | TACTTGTATCTTGAGTCGGCCATGATATACATGGCGG | 37 |
| 2 | TCACGAGCCAGCTCCGGCGGT | 21 | 14 | CTTTGCTGGCAGCATTCAGCTCGATGCGGTGTCGCCC | 37 |
| 3 | GTTGCCGTGTTGCCGGTGTTCTGCTGGTATCCAGCAG | 37 | 15 | CGAGAGTGATCCCGGGACCAGGATGGGCAC | 30 |
| 4 | TGAAGCACTGCACGCCGTAGGTACGTAGC | 29 | 16 | GACCATGTGATCGCGCTTCTCGCCGGACAC | 30 |
| 5 | GAAGAAGATGGTGCGCTCCTGGCAGGGTGG | 30 | 17 | GCTGAACTTGTGGCCGTTTACGTCGCCGTGGTGGGC | 36 |
| 6 | GAAGTCGATGCCCGCACGGGG | 21 | 18 | CCGTCCTTGGCTGTTGTAGTTG | 22 |
| 7 | GGTTGTCGCAGGGCGGGGTGGCAT | 24 | 19 | CACGAACGTGGTCGGCTTGTGCCCCAGGATCGTCCTT | 37 |
| 8 | ATGTGGTCGGGGCCCTTGCTCACCA | 25 | 20 | ATGAACTGCCCTCGTTGGGGT | 21 |
| 9 | CTTCGGGGACGTTGCGATGTTGTGGCGGACAGCTCGTCCATGC | 43 | 21 | CACCCCGGTGAACAGCTCCTCGTAGCGGC | 29 |
| 10 | TACTCCAGCGAGCTGCACGCTG | 22 | 22 | ACTTGAAGAAGTCGTCTGCTTAGTTCAC | 28 |
| 11 | CGCCCTCTCAGGGTCAGCTTGC | 22 | 23 | CAGGGCACGGGCAGCTTGCCGGTCTTGTA | 29 |
| 12 | CCGTCGCCGATGGGGTCCTCCTT | 23 | | | |

#### S1b. 3′ unhybridized — 23 staples

| # | Sequence (5′→3′) | nt | # | Sequence (5′→3′) | nt |
|---|---|---|---|---|---|
| 1 | GCTGAACTTGTGGCCGTTTACGTCGCCGTGGTGGGC | 36 | 13 | TGAAGCACTGCACGCCGTAGGTACGTAGC | 29 |
| 2 | TGGTTGTCTCAGGGCGGGTGGCAT | 24 | 14 | TACTCCAGGCGAGCTGCACGCT | 22 |
| 3 | GCCGTCCTGGCTGTTGTAGTTG | 22 | 15 | TCACGAGCCAGCTCGCGGCGG | 21 |
| 4 | CGCCCTCTCAGGGTCAGCTTGC | 22 | 16 | ATGAACTGCCCTCGGTTGGGG | 21 |
| 5 | GGACCATGTGATCGCGCTTCTCCCGGACAC | 30 | 17 | TCGAACTTCACCTCGGCGCGGGTGGTGCAG | 30 |
| 6 | TCACGAAAGTGGTCGCTTGTGCCCCAGGATCGTCCTT | 37 | 18 | CTTCGGGGACGTTGTCGATGTTGTGGCGG | 29 |
| 7 | ATCTTGAGTCGGCCATGATATACATGGCGG | 30 | 19 | GCCGTCGCCGATGGGTCCTCCTT | 23 |
| 8 | CACCCCGGTGAACAGCTCCTCGTAGCGGC | 29 | 20 | GAAGAAGATGGTGCGCTCCTGGCAGGGTGG | 30 |
| 9 | ATGTGGTCGGGGCCCTTGCTCACCA | 25 | 21 | TCTTTGCGGGCAGCTTCAGCTCGATGCGGTGTCGCCC | 37 |
| 10 | CCGAGAGTGATCCCGGACCAGGATGGGCAC | 30 | 22 | ACTTGAAGAAGTCGTCTGCTTAGTTCAC | 28 |
| 11 | GTTGCCGTGTTGCCGGGTGTTCTGCTGGTCTCCAGCA | 37 | 23 | CAGGGCACGGGCAGCTTGCCGGTCTTGTA | 29 |
| 12 | GAAGTCGATGCCCAGCACGGG | 21 | | | |

### S1c. 5′ unhybridized (35 nt coding overhang) — 21 staples

| # | Sequence (5′→3′) | nt | # | Sequence (5′→3′) | nt |
|---|---|---|---|---|---|
| 1 | CACGAACGTGGTCGGCTTGTGCCCCAGGATCGTCCTT | 37 | 12 | ACTTGAAGAAGTCGTCTGCTTAGTTCAC | 28 |
| 2 | TACTTGTATCTTGAGTCGGCCATGATATACATGGCGG | 37 | 13 | CCGTCCTTGGCTGTTGTAGTTG | 22 |
| 3 | CAGGGCACGGGCAGCTTGCCGGTCTTGTA | 29 | 14 | GCTGAACTTGTGGCCGTTTACGTCGCCGTGGTGGGC | 36 |
| 4 | TCGAACTTCACCTCGGCGCGGGTGGTGCAG | 30 | 15 | CCGTCGCCGATGGGGTCCTCCTT | 23 |
| 5 | CGCCCTCTCAGGGTCAGCTTGC | 22 | 16 | CTTTGCTGGCAGCATTCAGCTCGATGCGGTGTCGCCC | 37 |
| 6 | GACCATGTGATCGCGCTTCTCGCCGGACAC | 30 | 17 | CTTCGGGGACGTTGCGATGTTGTGGCGGACAGCTCGTCCATGC | 43 |
| 7 | TCACGAGCCAGCTCCGGCGGT | 21 | 18 | CGAGAGTGATCCCGGGACCAGGATGGGCAC | 30 |
| 8 | ATGAACTGCCCTCGTTGGGGT | 21 | 19 | TACTCCAGCGAGCTGCACGCTG | 22 |
| 9 | GAAGAAGATGGTGCGCTCCTGGCAGGGTGG | 30 | 20 | GTTGCCGTGTTGCCGGTGTTCTGCTGGTATCCAGCAG | 37 |
| 10 | GAAGTCGATGCCCGCACGGGG | 21 | 21 | GTCGGGGTAGCGGCTGAAGCACTGCACGCCGTAGGTACGTAGC | 43 |
| 11 | GGTTGTCGCAGGGCGGGGTGGCAT | 24 | | | |

### S1d. 5′ and 3′ unhybridized — 21 staples

| # | Sequence (5′→3′) | nt | # | Sequence (5′→3′) | nt |
|---|---|---|---|---|---|
| 1 | CAGGGCACGGGCAGCTTGCCGGTCTTGTA | 29 | 12 | TCGAACTTCACCTCGGCGCGGGTGGTGCAG | 30 |
| 2 | TCTTTGCGGGCAGCTTCAGCTCGATGCGGTGTCGCCC | 37 | 13 | ACTTGAAGAAGTCGTCTGCTTAGTTCAC | 28 |
| 3 | CGCCCTCTCAGGGTCAGCTTGC | 22 | 14 | TCACGAGCCAGCTCGCGGCGG | 21 |
| 4 | GGACCATGTGATCGCGCTTCTCCCGGACAC | 30 | 15 | GAAGTCGATGCCCAGCACGGG | 21 |
| 5 | TCACGAAAGTGGTCGCTTGTGCCCCAGGATCGTCCTT | 37 | 16 | TGGTTGTCTCAGGGCGGGTGGCAT | 24 |
| 6 | CCGAGAGTGATCCCGGACCAGGA | 23 | 17 | TACTCCAGGCGAGCTGCACGCT | 22 |
| 7 | GTTGCCGTGTTGCCGGGTGTTCTGCTGGTCTCCAGCA | 37 | 18 | GCCGTCGCCGATGGGTCCTCCTT | 23 |
| 8 | ATCTTGAGTCGGCCATGATATACATGGCGG | 30 | 19 | ATGAACTGCCCTCGGTTGGGG | 21 |
| 9 | CTTCGGGGACGTTGTCGATGTTGTGGCGG | 29 | 20 | TCGGGGTAGCGGCTGAAGCACTGCACGCCGTAGGTACGTAGC | 42 |
| 10 | GCCGTCCTGGCTGTTGTAGTTG | 22 | 21 | GAAGAAGATGGTGCGCTCCTGGCAGGGTGG | 30 |
| 11 | GCTGAACTTGTGGCCGTTTACGTCGCCGTGGTGGGC | 36 | | | |

### S1e. 50 nt coding overhang — 22 staples

| # | Sequence (5′→3′) | nt | # | Sequence (5′→3′) | nt |
|---|---|---|---|---|---|
| 1 | AGAGTGATCCCGGCGCCCTCGCCGGACACG | 30 | 12 | GGGCATGGCGGACTTGAAGAAGTCGGATGGTG | 32 |
| 2 | AGCGGCTGAAGCACTGCACGCCTTCACCT | 29 | 13 | GGGTGGTGTGGTGCGGGGTCT | 21 |
| 3 | AGTTGCCGTCGTCCTTGAAGAATGCTGCTT | 30 | 14 | GTCCTCGCATGATATAGACGTT | 22 |
| 4 | CACCAGGGTGTCGCCCTCGAACGTAGGTCA | 30 | 15 | GTCGCCGATGGGGGGCCCCAGGA | 23 |
| 5 | CATGTGGGCCCTCGGCGGTCA | 21 | 16 | GTGGCTGGAGCTGCACGCTGCC | 22 |
| 6 | CCATGTGATCGCGCTTCTCGTTAGATGAAC | 30 | 17 | TGTTGCCGTCCTCACGGGGCC | 21 |
| 7 | CGAACTCGGTCGGCTTGTAGTTGTACTCCGGTCTTGT | 37 | 18 | TTACTTGTACCTTGAAGTTCA | 21 |
| 8 | CGCTCCTTGTCGGCATGTTGTGGCGGATAGCTCGTCCATGCCG | 43 | 19 | TTCAGGGTCAGCTTGCCGTAGGTGGCATCTCGGGGT | 36 |
| 9 | CGGCGCGAGCTTGTTGTTCTGCTGGTAGTCAGCAGGA | 37 | 20 | TTGATGCCGTTCTTCTGCTGGACGTAGCCT | 30 |
| 10 | CTGAACTTGTGGCCGTTTACGTCGCCGTCCAGCTCG | 36 | 21 | TTGCTCACAGCAGCCTTGAAGTCGATGCCATGCGGTT | 37 |
| 11 | CTTGCCGCACGAGGGTGGGCCA | 22 | 22 | TTGTCGGGGGGCGGACACGGGCAG | 24 |

Constructs S1e and S1f differ by a single staple: CTGAACTTGTGGCCGTTTACGTCGCCGTCCAGCTCG (36 nt) in the 50 nt construct is truncated to CTGAACTTGTGGCCGTTTACGTCGCC (26 nt) in the 60 nt construct, extending the unhybridized region by 10 nt. The remaining 21 staples are identical.

### S1f. 60 nt coding overhang — 22 staples

| # | Sequence (5′→3′) | nt | # | Sequence (5′→3′) | nt |
|---|---|---|---|---|---|
| 1 | AGAGTGATCCCGGCGCCCTCGCCGGACACG | 30 | 12 | GGGCATGGCGGACTTGAAGAAGTCGGATGGTG | 32 |
| 2 | AGCGGCTGAAGCACTGCACGCCTTCACCT | 29 | 13 | GGGTGGTGTGGTGCGGGGTCT | 21 |
| 3 | AGTTGCCGTCGTCCTTGAAGAATGCTGCTT | 30 | 14 | GTCCTCGCATGATATAGACGTT | 22 |
| 4 | CACCAGGGTGTCGCCCTCGAACGTAGGTCA | 30 | 15 | GTCGCCGATGGGGGGCCCCAGGA | 23 |
| 5 | CATGTGGGCCCTCGGCGGTCA | 21 | 16 | GTGGCTGGAGCTGCACGCTGCC | 22 |
| 6 | CCATGTGATCGCGCTTCTCGTTAGATGAAC | 30 | 17 | TGTTGCCGTCCTCACGGGGCC | 21 |
| 7 | CGAACTCGGTCGGCTTGTAGTTGTACTCCGGTCTTGT | 37 | 18 | TTACTTGTACCTTGAAGTTCA | 21 |
| 8 | CGCTCCTTGTCGGCATGTTGTGGCGGATAGCTCGTCCATGCCG | 43 | 19 | TTCAGGGTCAGCTTGCCGTAGGTGGCATCTCGGGGT | 36 |
| 9 | CGGCGCGAGCTTGTTGTTCTGCTGGTAGTCAGCAGGA | 37 | 20 | TTGATGCCGTTCTTCTGCTGGACGTAGCCT | 30 |
| 10 | CTGAACTTGTGGCCGTTTACGTCGCC | 26 | 21 | TTGCTCACAGCAGCCTTGAAGTCGATGCCATGCGGTT | 37 |
| 11 | CTTGCCGCACGAGGGTGGGCCA | 22 | 22 | TTGTCGGGGGGCGGACACGGGCAG | 24 |

### S1g. 70 nt coding overhang — 22 staples

| # | Sequence (5′→3′) | nt | # | Sequence (5′→3′) | nt |
|---|---|---|---|---|---|
| 1 | AACTTCAGGGTCAGCTTGCCGTAGGTGGCTGGTCGG | 36 | 12 | GGTAGCGGCTGAAGCACTGCACAACTTCA | 29 |
| 2 | ACCTTGATGCCGTTCTTCTCCTGGACGTAG | 30 | 13 | GTGCGCTGCTTGTCTCGATGTTGTGGCGTACAGCTCGTCCATG | 43 |
| 3 | ACGCTGAACTTGTGGCCGT | 19 | 14 | GTTCACCAGGGTGTCGCCCTCGGCCGTAGG | 30 |
| 4 | CAGCTTGGGTCACGAGGGTGGG | 22 | 15 | GTTGTGGGGCGAGCTGCACGCT | 22 |
| 5 | CCGAGAGTGATCCCGTCGCCCTCGCCGGAC | 30 | 16 | TCACGAAAGTGGTCCTGTTGTAGTTGTACGCGGGTCT | 37 |
| 6 | CCTCGGCTCCAGCTGGGTGTTCTGCTGGTCTCCAGCA | 37 | 17 | TCAGGGTCCGGTGGGTTGGGG | 21 |
| 7 | CTTCATGATCGCCCGCGGCGG | 21 | 18 | TCTTTGCGGGCAGCCTCCTTGAAGTCGATTCGATGCG | 37 |
| 8 | GCCGTCCGGCCATGATATAGAC | 22 | 19 | TGGTTGTCTCAGGGCGGGCACGGG | 24 |
| 9 | GCCGTCGCCGATGGTGTGCCCCA | 23 | 20 | TGTAGTTGCCGTCGTCCTTGAATCGTGCTG | 30 |
| 10 | GGACCATGTGATCGCGCTTCTCTGCAGATG | 30 | 21 | TTACTTGGATCTTGAAGT | 18 |
| 11 | GGATGTTGCCGTCAGCACGGG | 21 | 22 | TTCGGGCATGGCGGACTTGAAGAAGGAAGATG | 32 |

#### S1h. 80 nt coding overhang — 21 staples

| # | Sequence (5′→3′) | nt | # | Sequence (5′→3′) | nt |
|---|---|---|---|---|---|
| 1 | GTGCGCTGCTTGTCTCGATGTTGTGGCGTACAGCTCGTCCATG | 43 | 12 | TGGTTGTCTCAGGGCGGGCACGGG | 24 |
| 2 | AACTTCAGGGTCAGCTTGCCGTAGGTGGCTGGTCGG | 36 | 13 | ACCTTGATGCCGTTCTTCTCCTGGACGTAG | 30 |
| 3 | CTTCATGATCGCCCGCGGCGG | 21 | 14 | GGATGTTGCCGTCAGCACGGG | 21 |
| 4 | TGTAGTTGCCGTCGTCCTTGAATCGTGCTG | 30 | 15 | TTCGGGCATGGCGGACTTGAAGAAGGAAGATG | 32 |
| 5 | CCGAGAGTGATCCCGTCGCCCTCGCCGGACACGCTGAAC | 39 | 16 | TCTTTGCGGGCAGCCTCCTTGAAGTCGATTCGATGCG | 37 |
| 6 | CCTCGGCTCCAGCTGGGTGTTCTGCTGGTCTCCAGCA | 37 | 17 | TCAGGGTCCGGTGGGTTGGGG | 21 |
| 7 | GTTGTGGGGCGAGCTGCACGCT | 22 | 18 | GGACCATGTGATCGCGCTTCTCTGCAGATG | 30 |
| 8 | TCACGAAAGTGGTCCTGTTGTAGTTGTACGCGGGTCT | 37 | 19 | GCCGTCGCCGATGGTGTGCCCCA | 23 |
| 9 | GCCGTCCGGCCATGATATAGAC | 22 | 20 | TTACTTGGATCTTGAAGT | 18 |
| 10 | CAGCTTGGGTCACGAGGGTGGG | 22 | 21 | GGTAGCGGCTGAAGCACTGCACAACTTCA | 29 |
| 11 | GTTCACCAGGGTGTCGCCCTCGGCCGTAGG | 30 | | | |

#### S1i. 100 nt coding overhang — 20 staples

| # | Sequence (5′→3′) | nt | # | Sequence (5′→3′) | nt |
|---|---|---|---|---|---|
| 1 | GCCGTTCGCTGGTAGTGGTCGG | 22 | 11 | TTCGGGCGATGAACGAGAGTG | 21 |
| 2 | ACGAACTCCAGCAGGACCATGTGGGTGGGC | 30 | 12 | TGTCGCCCTCGAACTTCACCTCACGTAGCC | 30 |
| 3 | TTACTTGTACAGCTCGTCCATGCCTTCAGGGTCAGCTTG | 39 | 13 | CTTGAAGAAGATGGTGCGCTCCTGGGGCGCGGGTCTTGTAGTTGCCGTCG | 50 |
| 4 | CTCCTTGAAGTCGATGCCCTTCCATGTGGT | 30 | 14 | GTTGTGGCGGATCTTCACGCTGCCGTCCTC | 30 |
| 5 | CGGGGTAGTCACGAGATCGCG | 21 | 15 | GGCTGTTGTAGTTGTTGTCGG | 21 |
| 6 | CAGGGCACGGGCAGCTTGCCGGTGGTGCAATGGCGG | 36 | 16 | CGAGCTGGAAGTTCACCTTGAT | 22 |
| 7 | ATCCCGGGTGTTCTTTCTGCTTGTCGGCCCACCAGGG | 37 | 17 | GGTGCTCATTGGGGTCCCGTAGGT | 24 |
| 8 | ACTTGAAGAAGTCGTGCTGCTTAGCTCGA | 29 | 18 | GCAGCAGCACGGGGAGACGTTGT | 23 |
| 9 | TGCGGTTATGATATCCGTCGCCGATGGGGCGGCGGTC | 37 | 19 | CTTCTCGGGTAGTGGTACTCCAGCTTGTGTTGCCGTC | 37 |
| 10 | CCGTAGGTGGCATCGCCCTCGC | 22 | 20 | CAGGGTGGCGGCTGAAGCACTG | 22 |

All staples were purchased from Integrated DNA Technologies, desalted, and used without further purification. Sequences are given 5′→3′. Staple lengths are stated for each strand; ranges and totals per construct are given in the summary table above.

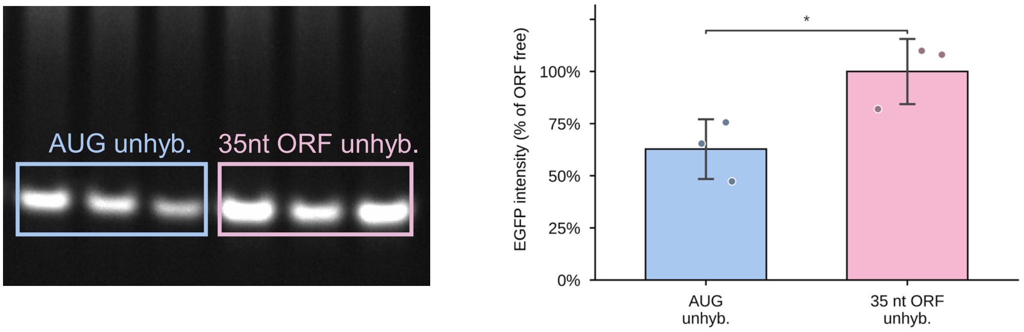


**Supplementary Figure S1. Exposing only the start codon gives lower translational output than exposing the first 35 coding nucleotides.** Left, native PAGE of the in vitro translation reactions, imaged in the Alexa 488 channel; in-gel fluorescence reports the folded EGFP product. Right, quantified EGFP fluorescence normalised to the 35 nt construct (= 100%). The construct leaving only the start codon unhybridized produced less output of the construct leaving the first 35 coding nucleotides unhybridized. *$p < 0.05$ (two-sample t-test). This comparison established 35 nt as the lower bound of the coding-overhang series, consistent with the 25–35 nt of accessible 5′ RNA required for efficient 43S loading.

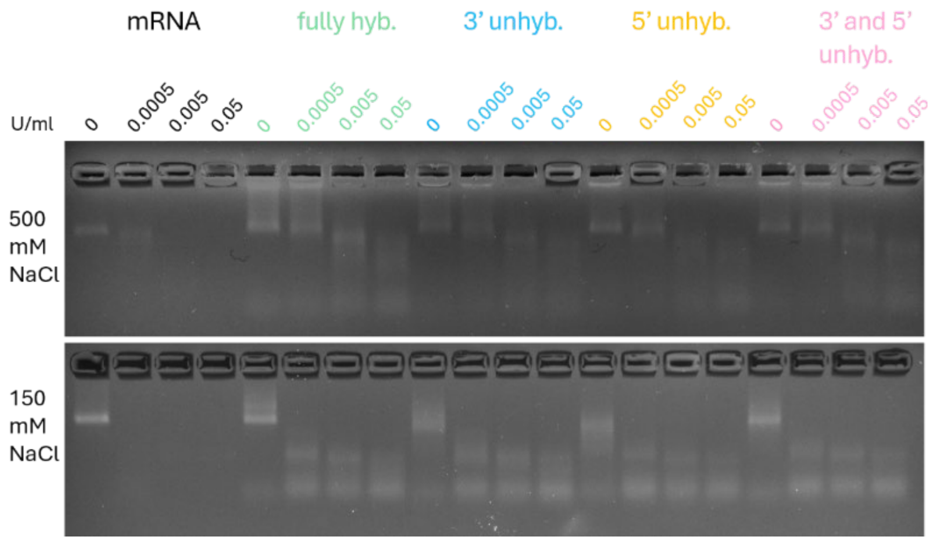


**Supplementary Figure S2. RNase A protection of the folded constructs.** Agarose gel electrophoresis of the fully hybridized, 3′ unhybridized, 5′ unhybridized, and 3′ and 5′ unhybridized constructs after incubation with RNase A at 0, 0.0005, 0.005, and 0.05 U ml$^{-1}$. The upper row shows digestion at 500 mM NaCl and the lower row at 150 mM NaCl. Within each construct group, lanes correspond to increasing RNase A concentration (left to right). Staple hybridization shields the mRNA scaffold from single-strand-specific nuclease digestion: the folded constructs retain a defined band at RNase A concentrations that degrade unprotected RNA, with the unhybridized single-stranded regions remaining the most susceptible to cleavage. Gels were run in 1× TAE with 11 mM $MgCl_2$ and stained with ethidium bromide

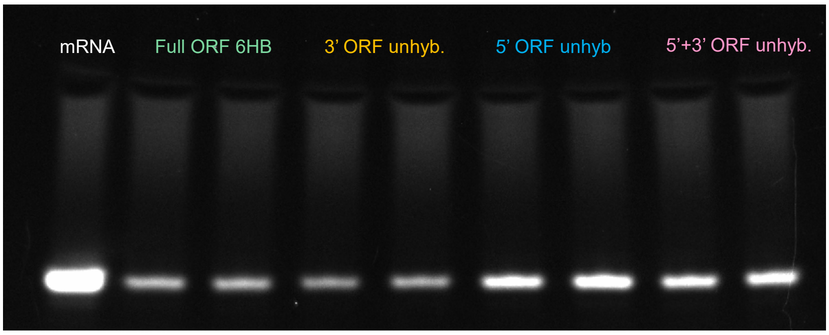


**Supplementary Figure S3.** Agarose gel electrophoresis (3.5% agarose, 1× TAE, 11 mM $MgCl_2$) of the four constructs in which a defined 35 nt region of the coding sequence is left unhybridized, alongside the naked mRNA scaffold. Left to right: mRNA scaffold; fully ORF-hybridized 6HB; 3′ ORF unhybridized; 5′ ORF unhybridized; 5′ and 3′ ORF unhybridized, each in duplicate. Folded species migrate as a compact band of lower mobility than the unhybridized scaffold, indicating successful assembly in all four designs. Gel stained with ethidium bromide and imaged under UV illumination.

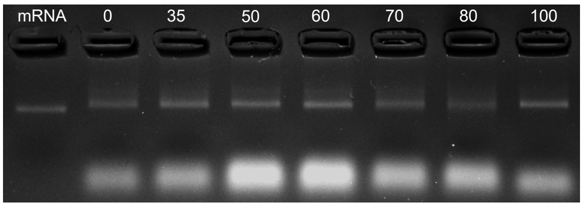


**Supplementary Figure S4. Folding of the overhang length-series constructs.** Agarose gel electrophoresis of the unmodified-uridine constructs across the 5′ coding overhang length series. The leftmost lane (mRNA) is the naked, unfolded mRNA scaffold; remaining lanes are folded origami with unstapled 5′ overhangs of 0 (fully hybridized), 35, 50, 60, 70, 80, and 100 nt. All constructs migrate as a compact, retarded band relative to the scaffold, confirming successful folding across the series, with excess staple strands running ahead as the fast-migrating lower band. Gels were run in 1× TAE with 11 mM $MgCl_2$ and stained with ethidium bromide.

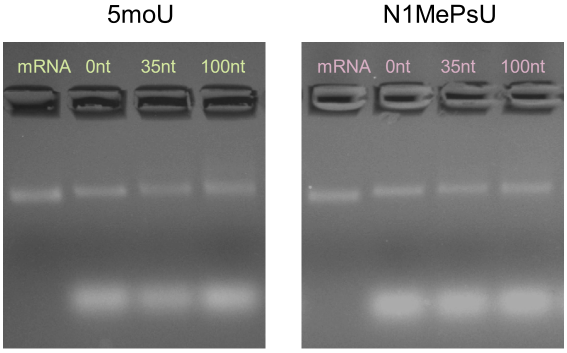


**Supplementary Figure S5. Folding of the modified-uridine constructs.** Agarose gel electrophoresis of the 5-methoxyuridine (5moU, left) and N1-methylpseudouridine (N1MePsU, right) constructs at 0 (fully hybridized), 35, and 100 nt overhang length. The leftmost lane of each gel (mRNA) is the corresponding naked, unfolded modified mRNA scaffold; remaining lanes are folded origami at the indicated overhang length. All modified constructs migrate as a compact, lower mobility band relative to the scaffold, confirming that 5moU and N1MePsU substitution does not impair folding of the hybrid origami. Gels were run in 1× TAE with 11 mM $MgCl_2$ and stained with ethidium bromide.

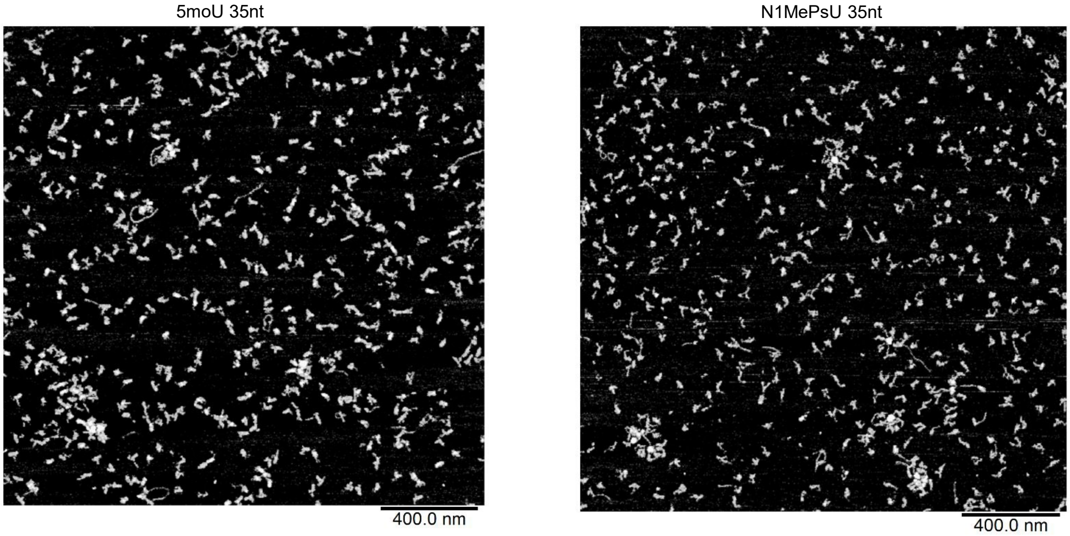


**Supplementary Figure S6. AFM of the modified-uridine constructs.** Representative atomic force microscopy images of the 35 nt overhang constructs folded with 5-methoxyuridine (5moU, left) and N1-methylpseudouridine (N1MePsU, right). Both modified constructs form rod-shaped particles comparable to the unmodified-uridine constructs (Figure 2E), confirming that 5moU and N1MePsU substitution does not alter the folded morphology of the hybrid origami. Scale bars, 400 nm.

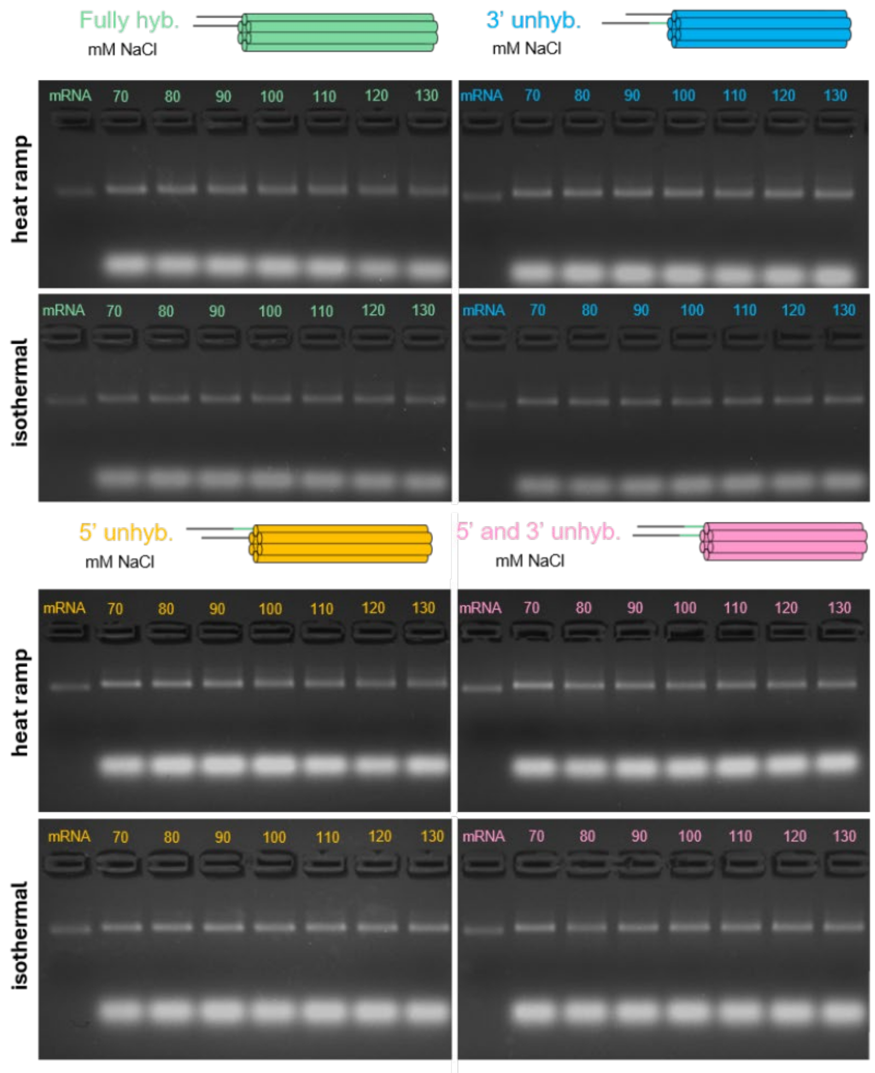


**Supplementary Figure S7. NaCl concentration screen for folding of the four position-experiment constructs.** Agarose gel electrophoresis of the fully hybridized, 3′ unhybridized, 5′ unhybridized, and 5′ and 3′ unhybridized constructs folded across NaCl concentrations from 70 to 130 mM (in 10 mM Tris). For each construct, two independent folding reactions are shown (upper and lower rows). The leftmost lane of each gel (mRNA) is the naked, unfolded mRNA scaffold; remaining lanes are folded origami at the indicated NaCl concentration (mM). Folded structures migrate as a compact, lower mobility band relative to the scaffold, with excess staple strands running ahead as the fast-migrating lower band. Folding was efficient across the full range screened; 110 mM NaCl was selected for all subsequent experiments. Gels were run in 1× TAE with 11 mM $MgCl_2$ and stained with ethidium bromide.

### Supplementary Table S2. Position of the unhybridized coding region

#### S2a. Individual replicate values

| Condition | Rep 1 | Rep 2 | Rep 3 | Rep 4 | Rep 5 | Mean (AU) | SD (AU) |
|---|---|---|---|---|---|---|---|
| Fully ORF-hybridized | 8338.3 | 7889.3 | 6666.6 | 11445.8 | 10579.0 | 8983.8 | 1974.2 |
| 3′ coding region unstapled | 6114.8 | 7916.1 | 5177.7 | 2855.9 | 7424.6 | 5897.8 | 2013.9 |
| 5′ coding region unstapled (ref) | 13187.6 | 14489.1 | 11037.8 | 16169.3 | 14217.9 | 13820.3 | 1888.5 |
| 5′ + 3′ coding regions unstapled | 11115.8 | 11360.7 | 14158.0 | 12696.0 | 12183.8 | 12302.9 | 1215.2 |

AU = arbitrary fluorescence units. n = 5 independent folding reactions and independent translation reactions per condition. One-way ANOVA $F(3,16) = 19.24$, $p < 0.0001$. Reference = 5′ coding region unstapled.

#### S2b. Summary statistics, normalised to the 5′-free reference

| Condition | Mean (AU) | SD (AU) | Norm. (% of 5′-free) | SD (%) | Tukey vs ref |
|---|---|---|---|---|---|
| Fully ORF-hybridized | 8983.8 | 1974.2 | 65.0% | 14.3% | ** (p = 0.0031) |
| 3′ coding region unstapled | 5897.8 | 2013.9 | 42.7% | 14.6% | *** (p < 0.0001) |
| 5′ coding region unstapled (ref) | 13820.3 | 1888.5 | 100.0% | 13.7% | ref |
| 5′ + 3′ coding regions unstapled | 12302.9 | 1215.2 | 89.0% | 8.8% | ns (p = 0.558) |

Normalised to the 5′ coding region unstapled construct (= 100%). Remaining pairwise comparisons (Tukey HSD): 3′ vs fully ORF-hybridized $p = 0.067$; 3′ vs 5′+3′ $p = 0.0002$; fully ORF-hybridized vs 5′+3′ $p = 0.045$. Variances were homogeneous (Levene $p = 0.822$; Bartlett $p = 0.79$) and all conditions were consistent with normality (Shapiro–Wilk $p \geq 0.599$). ** $p < 0.01$, *** $p < 0.001$, ns = not significant.

### Supplementary Table S3. 5′ coding overhang length series

#### S3a. Individual replicate values

| nt | Rep 1 | Rep 2 | Rep 3 | Rep 4 | Rep 5 | Rep 6 | Mean (AU) | SD (AU) |
|---|---|---|---|---|---|---|---|---|
| 35 | 19113.5 | 15329.8 | 20478.0 | 19338.8 | 15278.2 | 20304.3 | 18307.1 | 2385.6 |
| 50 | 9683.8 | 11778.4 | 18235.2 | 18379.3 | 10659.0 | 10659.1 | 13232.5 | 3986.7 |
| 60 | 9484.6 | 15138.2 | 14462.3 | 15881.9 | 13452.5 | 9750.7 | 13028.4 | 2761.6 |
| 70 | 8552.8 | 12957.6 | 9778.2 | 13054.7 | 9121.5 | 9112.4 | 10429.5 | 2033.5 |
| 80 | 11914.2 | 16640.9 | 13485.2 | 16784.4 | 12438.0 | 12817.9 | 14013.4 | 2153.1 |
| 100 | 4501.4 | 8261.6 | 6836.1 | 6133.6 | 4866.2 | 8599.4 | 6533.0 | 1697.9 |

AU = arbitrary fluorescence units. n = 6 independent folding reactions and independent translation reactions per condition. One-way ANOVA $F(5,30) = 13.51$, $p < 0.0001$ ($MSE = 6.811 \times 10^6\ AU^2$). Reference = 35 nt construct.

#### S3b. Summary statistics, normalised to the 35 nt reference

| Overhang (nt) | Mean (AU) | SD (AU) | Norm. (% of 35 nt) | SD (%) | Tukey vs 35 nt |
|---|---|---|---|---|---|
| 35 | 18307.1 | 2385.6 | 100.0% | 13.0% | ref |
| 50 | 13232.5 | 3986.7 | 72.3% | 21.8% | * (p = 0.023) |
| 60 | 13028.4 | 2761.6 | 71.2% | 15.1% | * (p = 0.017) |
| 70 | 10429.5 | 2033.5 | 57.0% | 11.1% | *** (p = 0.0002) |
| 80 | 14013.4 | 2153.1 | 76.5% | 11.8% | ns (p = 0.077) |
| 100 | 6533.0 | 1697.9 | 35.7% | 9.3% | *** (p < 0.0001) |

Normalised to the 35 nt construct (= 100%). Comparisons in this column are against the 35 nt reference only; the complete pairwise matrix is given in Supplementary Table S3c and should be consulted before interpreting any construct relative to its neighbours. Variances were homogeneous (Levene p = 0.767; Bartlett p = 0.49). Shapiro–Wilk indicated departures from normality at 50 nt (p = 0.030) and 70 nt (p = 0.044); ANOVA and Tukey HSD are robust to departures of this magnitude at equal n. * p < 0.05, *** p < 0.001, ns = not significant.

#### S3c. Complete pairwise comparison matrix (Tukey HSD, all 15 pairs)

| Pair (nt) | Δ (AU) | Δ (% of 35 nt) | q | p (Tukey) | Significance |
|---|---|---|---|---|---|
| 35 vs 50 | 5075 | 27.7 | 4.76 | 0.023 | * |
| 35 vs 60 | 5279 | 28.8 | 4.95 | 0.017 | * |
| 35 vs 70 | 7878 | 43.0 | 7.39 | 0.0002 | *** |
| 35 vs 80 | 4294 | 23.5 | 4.03 | 0.077 | ns |
| 35 vs 100 | 11774 | 64.3 | 11.05 | < 0.0001 | *** |
| 50 vs 60 | 204 | 1.1 | 0.19 | 1.000 | ns |
| 50 vs 70 | 2803 | 15.3 | 2.63 | 0.445 | ns |
| 50 vs 80 | 781 | 4.3 | 0.73 | 0.995 | ns |
| 50 vs 100 | 6699 | 36.6 | 6.29 | 0.0014 | ** |
| 60 vs 70 | 2599 | 14.2 | 2.44 | 0.527 | ns |
| 60 vs 80 | 985 | 5.4 | 0.92 | 0.986 | ns |
| 60 vs 100 | 6495 | 35.5 | 6.10 | 0.0020 | ** |
| 70 vs 80 | 3584 | 19.6 | 3.36 | 0.196 | ns |
| 70 vs 100 | 3896 | 21.3 | 3.66 | 0.132 | ns |
| 80 vs 100 | 7480 | 40.9 | 7.02 | 0.0003 | *** |

All 15 pairwise comparisons among the six overhang lengths. $\sqrt{(MSE/n)}$ = 1065 AU; $q^{crit}(6, 30, \alpha = 0.05) = 4.301$, giving an honestly significant difference of 4583 AU: any two means differing by less than this are not distinguishable at α = 0.05. The 80 nt construct is not distinguishable from the 50, 60 or 70 nt constructs (p = 0.995, 0.986 and 0.196 respectively). Only the 100 nt construct is clearly separated from the interior of the series. * p < 0.05, ** p < 0.01, *** p < 0.001, ns = not significant.

### Supplementary Table S3d. Regime analysis of the 5′ coding overhang length series

#### S3d(i). Output grouped by regime

| Regime | n | Mean (AU) | SD (AU) | Norm. (% of 35 nt) | Tukey (all pairs) |
|---|---|---|---|---|---|
| 35 nt | 6 | 18307.1 | 2385.6 | 100.0% | — |
| Interior, 50–80 nt pooled | 24 | 12675.9 | 2986.0 | 69.2% | vs 35 nt: p = 0.0002 |
| 100 nt | 6 | 6533.0 | 1697.9 | 35.7% | vs interior: p = 0.0001 |

One-way ANOVA across the three regimes: $F(2,33) = 27.71$, $p < 0.0001$. All three pairwise comparisons are significant (Tukey HSD): 35 nt vs interior, Δ = 5631 AU [2561, 8701], p = 0.0002; interior vs 100 nt, Δ = 6143 AU [3073, 9213], p = 0.0001; 35 nt vs 100 nt, Δ = 11774 AU [7891, 15657], p < 0.0001. Square brackets give 95% confidence intervals on the difference in means.

#### S3d(ii). Homogeneity within the interior regime

| Overhang (nt) | Mean (AU) | SD (AU) | Norm. (% of 35 nt) | ANOVA across 50–80 nt |
|---|---|---|---|---|
| 50 | 13232.5 | 3986.7 | 72.3% | F(3,20) = 1.80 |
| 60 | 13028.4 | 2761.6 | 71.2% | p = 0.179 |
| 70 | 10429.5 | 2033.5 | 57.0% | not significant |
| 80 | 14013.4 | 2153.1 | 76.5% | — |

Translational output does not differ among the 50, 60, 70 and 80 nt constructs. A Jonckheere–Terpstra test for monotone trend restricted to these four constructs is likewise non-significant (z = 0.05, p = 0.959). Output within this interval is therefore independent of overhang length, and the 80 nt construct is not distinguishable from the other members of the interior regime.

#### S3d(iii). Model comparison on construct means

| Model | $R^2$ | AIC | Description |
|---|---|---|---|
| Linear in overhang length | 0.737 | 111.20 | Output declines continuously with length |
| Three-regime step | 0.905 | 107.07 | 35 nt > interior (50–80 nt) > 100 nt |

The three-regime description fits the construct means better than a continuous decline (ΔAIC = 4.1). Output falls in two discrete steps rather than in proportion to overhang length: exposure beyond the ribosomal initiation footprint incurs a fixed penalty that does not increase between 50 and 80 nt, and a further penalty at 100 nt. The complete pairwise matrix for the six individual constructs is given in Supplementary Table S3c.

### Supplementary Table S4. RNAfold metrics and translational output

| Overhang (nt) | ΔG (kcal/mol) | MFE freq (%) | Ensemble div. | AUG accessibility | EGFP (% of 35 nt) |
|---|---|---|---|---|---|
| 35 | −17.10 | 13.9 | 6.73 | 0.307 | 100.0 |
| 50 | −26.00 | 5.7 | 8.27 | 0.287 | 72.3 |
| 60 | −28.50 | 6.9 | 12.22 | 0.289 | 71.2 |
| 70 | −32.40 | 7.3 | 9.16 | 0.289 | 57.0 |
| 80 | −34.50 | 2.1 | 14.75 | 0.290 | 76.5 |
| 100 | −49.50 | 7.7 | 5.38 | 0.286 | 35.7 |

ΔG, MFE frequency and ensemble diversity from the ViennaRNA partition function (ViennaRNA 2.7.2, Turner 2004 parameters, 37 °C). AUG accessibility = mean unpaired probability of the start codon and the first five codons (positions 49–63), from the full-construct partition function. EGFP output normalised to the 35 nt construct (= 100%), from Supplementary Table S3b.

### Supplementary Table S5. Correlation of RNAfold metrics with translational output (n = 6)

| RNAfold metric | Pearson r | $R^2$ | p | Interpretation |
|---|---|---|---|---|
| ΔG (minimum free energy) | +0.920 | 0.846 | 0.009 | Strongest correlate; co-varies with overhang length |
| MFE frequency | +0.356 | 0.127 | 0.488 | Not significant |
| Ensemble diversity | +0.258 | 0.067 | 0.621 | Not significant |
| AUG-region accessibility | +0.792 | 0.628 | 0.060 | Not significant; driven by the 35 nt construct alone |
| Overhang length | −0.859 | 0.738 | 0.028 | Co-varies with ΔG (r = −0.981, p = 0.0005) |

Pearson correlation with EGFP output (% of 35 nt) across the six constructs. ΔG is the only metric correlating with output at $p < 0.05$ on the full series, but it cannot be interpreted independently of overhang length, with which it co-varies almost perfectly. AUG-region accessibility is effectively constant across the 50–100 nt constructs (0.286–0.290); its apparent correlation reflects only the shorter 35 nt construct and does not reach significance.

#### S5b. Leave-one-out sensitivity of the ΔG correlation

| Construct omitted | Pearson r | $R^2$ | p | Effect |
|---|---|---|---|---|
| None (n = 6) | +0.920 | 0.846 | 0.009 | Full series |
| 35 nt | +0.857 | 0.735 | 0.063 | No longer significant |
| 50 nt | +0.931 | 0.866 | 0.022 | Retained |
| 60 nt | +0.922 | 0.849 | 0.026 | Retained |
| 70 nt | +0.943 | 0.889 | 0.016 | Retained |
| 80 nt | +0.970 | 0.942 | 0.006 | Retained |
| 100 nt | +0.798 | 0.637 | 0.105 | No longer significant |

The correlation is not robust to removal of either endpoint. Omitting the 100 nt construct, which lies far from the remaining five in both ΔG and output, reduces the correlation to r = +0.798 (p = 0.105); among the 35–80 nt constructs alone the relationship is not significant. The reported r = +0.920 on the full series is therefore leverage-dependent and should not be presented as an independent structural determinant of translational output.

### Supplementary Table S6. RNAfold MFE secondary structures (dot-bracket notation)

| nt | Seq len | MFE secondary structure (dot-bracket notation), unmodified uridine |
|---|---|---|
| 35 | 83 | ..........................................(((.((..((((((((...((.....))))))))))))))) |
| 50 | 98 | ..........................................((((((.(((((((((...((.....))))))))))).))))))(((......))) |
| 60 | 108 | .......................(((..((............((((((.(((((((((...((.....))))))))))).)))))).............))..))).. |
| 70 | 118 | ..........................................((((((.(((((((((...((.....))))))))))).))))))(((..(((.(......).))).)))...... |
| 80 | 128 | ..........................................((((((.(((((((((...((.....))))))))))).))))))........((((((......(((....)))...))))))... |
| 100 | 148 | ..........................................((((((.(((((((((...((.....))))))))))).))))))..((.((((((.((.((((((((((..........)))....)))<br>)))).)).)))).)))) |

Minimum-free-energy structures predicted by RNAfold (ViennaRNA 2.7.2). The first ~42 positions, shown unpaired, correspond to the 5′ UTR. These are the single lowest-energy folds; ensemble-level domain statistics are given in Supplementary Table S7. Sequence length is the 48 nt 5′ UTR plus the stated coding overhang

### Supplementary Table S7. Boltzmann ensemble domain-count distribution

| Overhang (nt) | MFE domains | Mean domains | 1 domain (%) | 2 domains (%) | ≥3 domains (%) | EGFP (% of 35 nt) |
|---|---|---|---|---|---|---|
| 35 | 1 | 1.12 | 88 | 12 | 0 | 100.0 |
| 50 | 2 | 1.59 | 45 | 51 | 4 | 72.3 |
| 60 | 1 | 1.63 | 41 | 55 | 5 | 71.2 |
| 70 | 2 | 2.02 | 6 | 86 | 8 | 57.0 |
| 80 | 2 | 2.06 | 5 | 84 | 11 | 76.5 |
| 100 | 2 | 2.07 | 0 | 93 | 7 | 35.7 |

5000 structures stochastically sampled from the Boltzmann ensemble per construct (uniq_ML = 1). Percentages are rounded to the nearest integer and may not sum to 100 (60 nt sums to 101). Domain count is ensemble-robust but does not track translational output: the 70 and 80 nt constructs have near-identical distributions yet differ in measured output, and the 80 and 100 nt constructs are both predominantly two-domain yet translate at 76.5% and 35.7% of the 35 nt reference.

### Supplementary Table S8. Correlation of ensemble domain metrics with translational output (n = 6)

| Ensemble metric | Pearson r | $R^2$ | p | Interpretation |
|---|---|---|---|---|
| Mean domain count | −0.793 | 0.628 | 0.060 | Not significant; scales with overhang length |
| % 2-domain | −0.843 | 0.710 | 0.035 | Scales with overhang length |
| % 1-domain | +0.816 | 0.666 | 0.048 | Scales with overhang length |

These correlations reflect co-variation of domain count with overhang length rather than an independent structural effect, and provide no explanation for the translational pattern beyond that already captured by overhang length and ΔG.

### Supplementary Table S9. 5moU overhang length series (0, 35, 100 nt)

#### S9a. Individual replicates

| Condition | Rep 1 | Rep 2 | Rep 3 | Rep 4 | Rep 5 | Rep 6 | Mean (AU) |
|---|---|---|---|---|---|---|---|
| 5moU 0 nt | 15521.409 | 15401.936 | 15473.284 | 12237.418 | 12659.774 | 12643.271 | 13989.5 |
| 5moU 35 nt | 20190.894 | 15073.187 | 19145.480 | 16814.203 | 19733.516 | 17871.959 | 18138.2 |
| 5moU 100 nt | 13900.702 | 7223.631 | 5194.617 | 6412.885 | 10847.092 | 8835.334 | 8735.7 |

AU = arbitrary fluorescence units. n = 6 independent folding reactions and independent translation reactions per condition. Values are reported at the precision of the ImageJ output.

#### S9b. Summary, normalised to 5moU 35 nt

| Condition | Mean (AU) | SD (AU) | Norm. (% of 35 nt) | SD (%) | CV (%) | Tukey vs 35 nt |
|---|---|---|---|---|---|---|
| 5moU 0 nt | 13989.5 | 1624.4 | 77.1% | 9.0% | 11.6% | * |
| 5moU 35 nt | 18138.2 | 1948.9 | 100.0% | 10.7% | 10.7% | ref |
| 5moU 100 nt | 8735.7 | 3205.2 | 48.2% | 17.7% | 36.7% | *** |

0 nt = fully ORF-hybridized. One-way ANOVA $F(2,15) = 23.92$, $p < 0.0001$. Pairwise (Tukey HSD): 0 vs 35 $p = 0.021$; 35 vs 100 $p < 0.0001$; 0 vs 100 $p = 0.0042$. Variances were homogeneous across conditions (Levene $p = 0.346$; Bartlett $p = 0.30$), so Tukey HSD is appropriate for this experiment. Shapiro–Wilk indicated a departure from normality for the 0 nt condition ($p = 0.020$); a Games–Howell sensitivity analysis returns the same qualitative outcome (0 vs 35 $p = 0.007$; 35 vs 100 $p = 0.001$; 0 vs 100 $p = 0.020$). * $p < 0.05$, *** $p < 0.001$.

### Supplementary Table S10. N1MePsU overhang length series (0, 35, 100 nt)

#### S10a. Individual replicates

| Condition | Rep 1 | Rep 2 | Rep 3 | Rep 4 | Rep 5 | Rep 6 | Mean (AU) |
|---|---|---|---|---|---|---|---|
| N1MePsU 0 nt | 12909.9 | 12345.1 | 12052.7 | 12436.5 | 12566.1 | 12316.5 | 12437.8 |
| N1MePsU 35 nt | 14793.6 | 15376.1 | 15274.5 | 14871.0 | 15006.6 | 14839.5 | 15026.9 |
| N1MePsU 100 nt | 1870.9 | 2352.7 | 2203.7 | 2480.9 | 2844.0 | 2324.2 | 2346.1 |

AU = arbitrary fluorescence units. n = 6 independent folding reactions and independent translation reactions per condition.

#### S10b. Summary, normalised to N1MePsU 35 nt

| Condition | Mean (AU) | SD (AU) | Norm. (% of 35 nt) | SD (%) | CV (%) | Tukey vs 35 nt |
|---|---|---|---|---|---|---|
| N1MePsU 0 nt | 12437.8 | 286.7 | 82.8% | 1.9% | 2.3% | *** |
| N1MePsU 35 nt | 15026.9 | 243.9 | 100.0% | 1.6% | 1.6% | ref |
| N1MePsU 100 nt | 2346.1 | 320.2 | 15.6% | 2.1% | 13.6% | *** |

0 nt = fully ORF-hybridized. One-way ANOVA $F(2,15) = 3308.6$, $p < 0.0001$. All three pairwise comparisons $p < 0.0001$ (Tukey HSD). Variances were homogeneous (Levene $p = 0.982$; Bartlett $p = 0.85$). *** $p < 0.001$.

### Supplementary Table S11. Cross-modification comparison at fixed 35 nt overhang

#### S11a. Individual replicates

| Condition | Rep 1 | Rep 2 | Rep 3 | Rep 4 | Rep 5 | Rep 6 | Mean (AU) |
|---|---|---|---|---|---|---|---|
| Uridine (U) | 9467.1 | 11273.1 | 15732.9 | 13759.7 | 11711.8 | 14246.6 | 12698.5 |
| 5moU | 15020.3 | 18641.2 | 13985.5 | 19605.5 | 15309.9 | 15309.9 | 16312.0 |
| N1MePsU | 20164.6 | 18267.5 | 19526.3 | 20843.9 | 20060.3 | 18866.3 | 19621.5 |

AU = arbitrary fluorescence units. n = 6 independent folding reactions and independent translation reactions per condition. All three conditions share an identical 35 nt free 5′ coding overhang; only the scaffold uridine chemistry differs.

#### S11b. Summary, normalised to N1MePsU

| Condition | Mean (AU) | SD (AU) | Norm. (% of N1MePsU) | CV (%) | Tukey |
|---|---|---|---|---|---|
| Uridine (U) | 12698.5 | 2288.1 | 64.7% | 18.0% | *** vs N1MePsU; * vs 5moU |
| 5moU | 16312.0 | 2252.2 | 83.1% | 13.8% | * vs N1MePsU; * vs U |
| N1MePsU (ref) | 19621.5 | 937.5 | 100.0% | 4.8% | ref |

One-way ANOVA $F(2,15) = 19.29$, $p < 0.0001$. Pairwise (Tukey HSD): U vs 5moU $p = 0.014$; U vs N1MePsU $p < 0.0001$; 5moU vs N1MePsU $p = 0.025$. Variances were homogeneous (Levene $p = 0.294$; Bartlett $p = 0.16$). * $p < 0.05$, *** $p < 0.001$. CV = coefficient of variation.

### Supplementary Table S11c. Chemistry × overhang analysis (5moU and N1MeΨ)

#### S11c(i). Two-way ANOVA on log-transformed output

| Term | df | F | p | Interpretation |
|---|---|---|---|---|
| Uridine chemistry | 1, 30 | 83.76 | < 0.0001 | Chemistries differ in absolute output |
| Overhang length | 2, 30 | 205.87 | < 0.0001 | Output depends on overhang |
| Chemistry × overhang | 2, 30 | 43.14 | < 0.0001 | Response differs between chemistries |

Computed from the replicate values in Supplementary Tables S9a and S10a. Output was log-transformed so that the interaction term tests whether the proportional (fold-change) response to overhang length differs between chemistries, independently of any difference in absolute signal between experiments.

#### S11c(ii). Interaction decomposed by contrast

| Contrast | F (1, 20) | p | 5moU | N1MeΨ | Ratio of |
|---|---|---|---|---|---|
| 0 vs 35 nt | 1.14 | 0.299 | 77.1% | 82.8% | 0 / 35 |
| 35 vs 100 nt | 43.87 | < 0.0001 | 45.8% | 15.5% | 100 / 35 |
| 0 vs 100 nt | 49.33 | < 0.0001 | 59.5% | 18.7% | 100 / 0 |

Each row tests whether the proportional change between two overhang conditions differs between the chemistries. Percentages are geometric mean ratios between the two conditions named in the final column, computed within each chemistry. The benefit of exposing the proximal coding region (0 vs 35 nt) is quantitatively indistinguishable between the two chemistries, whereas the penalty for exposing 100 nt is not. The third row uses a contrast that does not involve the 35 nt construct and reproduces the same result, establishing that the difference resides in the 100 nt condition itself rather than in normalisation to each chemistry's own reference: if the N1MeΨ 35 nt value were disproportionately high, its 0 nt ratio would also be depressed, whereas that ratio is in fact higher than for 5moU. Sensitivity to an over-long overhang is approximately threefold greater for N1MeΨ.

**S11c(iii). Condition means underlying the contrasts**

| Chemistry | 0 nt (AU) | 35 nt (AU) | 100 nt (AU) | 100 nt as % of 0 nt |
|---|---|---|---|---|
| 5moU | 13989.5 | 18138.2 | 8735.7 | 59.5% |
| N1MeΨ | 12437.8 | 15026.9 | 2346.1 | 18.7% |

Arithmetic means from Supplementary Tables S9a and S10a. The two chemistries produce comparable absolute output at 0 and 35 nt but differ by almost fourfold at 100 nt, which is where the interaction reported above resides. Because these values derive from separate experiments, absolute intensities are not directly comparable between chemistries; the analyses in S11c(i) and S11c(ii) are performed on log-transformed data so that any constant scaling difference between experiments is absorbed into the chemistry main effect and does not contribute to the interaction term.
